# Inducing an oxidized redox-balance improves anti-tumor CD8^+^ T cell function

**DOI:** 10.1101/2023.03.27.533229

**Authors:** Ju Hee Oh, Rachel A. Cederberg, Erin Tanaka, Luisa Bopp, Terri Ser, Sara Niyyati, Annette E. Patterson, Neeku Amanat, Jared Dutra, Patricia Ye, Meredith Clark, Kirsten Ward-Hartstonge, Anne-Sophie Archambault, Janice Tsui, Philipp F. Lange, Sue Tsai, C. Bruce Verchere, Yongjin Park, Mario Fabri, Kevin L. Bennewith, Ramon I. Klein Geltink

**Affiliations:** BC Children’s Hospital Research Institute; Vancouver, BC, Canada; Department of Pathology and Laboratory Medicine, University of British Columbia; Vancouver, BC, Canada; BC Cancer Research Institute; Vancouver, BC, Canada; Department of Dermatology and Venereology, University of Cologne, Faculty of Medicine, and University Hospital of Cologne; Cologne, Germany; Interdisciplinary Oncology Program, University of British Columbia; Vancouver, BC, Canada; Department of Medical Microbiology and Immunology, University of Alberta; Edmonton, Canada; Center for Molecular Medicine Cologne (CMMC), University of Cologne; Cologne, Germany

**Keywords:** adoptive cellular therapy, CD8+ T cells, redox homeostasis, mitochondrial reactive oxygen species, immunotherapy of cancer, human tumor infiltrating lymphocytes.

## Abstract

Cancer immunotherapy using antigen-specific CD8^+^ T cells depends on long-lasting anti-tumor function of the *in vitro* expanded T cells. T cell function is intricately linked to the activity of many metabolic pathways directly impacting the ability of CD8^+^ T cells to kill tumor cells. Metabolic conditioning *in vitro* better prepares CD8^+^ T cells for *in vivo* survival, tumor infiltration and tumor clearance. The mechanism underlying *in vitro* metabolic conditioning-induced augmented *in vivo* T cell function remains poorly understood. Here we show that metabolic conditioning of CD8^+^ effector T cells induces an oxidized cellular redox balance at least in part mediated by increased mitochondrial reactive oxygen species (ROS). This redox shift contributes to enhanced *in vivo* persistence and tumor clearance. In human tumour-infiltrating T cells, altering the redox balance *ex-vivo* reinvigorated pro-inflammatory cytokine production. Therefore, we believe that redox alterations present a targetable pathway to increase T cell-based anti-tumor immunotherapy efficacy.

## INTRODUCTION

Cancer immunotherapy using tumor antigen-specific CD8^+^ T cells is a promising strategy, leading to complete remission in blood cancers ^1, 2^. However, efficacy in the context of solid tumors is disappointing, largely because the hostile tumor microenvironment (TME) limits the anti-tumor activity of these cells ^3–5^. Key roadblocks are: lack of tumor infiltration, poor survival and/or exhaustion of the immune cells that do infiltrate ^5, 6^, and the fact that CD8^+^ T cells and tumor cells have overlapping metabolic needs, leading to competition for nutrients within the TME ^4, 7, 8^. To kill cancer cells, CD8^+^ T cells activate a variety of metabolic pathways, such as glycolysis, mitochondrial metabolism, and the pentose phosphate pathway (PPP) often referred to as metabolic reprogramming. Mouse models of solid tumors show impaired mitochondrial metabolism in CD4^+^ and CD8^+^ T cells, dampening anti-tumor function ^9,10^. In humans, T cells isolated from tumors show mitochondrial dysfunction ^9^, and patients responding to immune checkpoint therapies that block inhibitory signaling receptors expressed by T cells ^11–13^ show increased glycolysis *and* mitochondrial metabolism in tumor-infiltrating CD8^+^ T cells ^14, 15^.

Together, these findings support the concept that increasing or sustaining both glycolytic and mitochondrial metabolism in CD8^+^ T cells could improve anti-tumor T cell function. Thus, strategies to maintain the beneficial metabolism and function of therapeutic T cell products for cellular therapy of solid cancer are crucial to improve outcomes. Restricting glucose metabolism during initial activation *in vitro* ^16^, selecting T cells with low mitochondrial membrane potential ^17^, glucose restriction of fully activated CD8^+^ T cells ^18^, genetic or pharmacological inhibition of the PPP ^19^, and exposure of CD8^+^ T cells to increased concentrations of lactate ^20^ or potassium ^21^ are some examples of metabolic strategies to improve the efficacy of anti-tumor T cell function. It was hypothesized that these strategies induce memory precursor or stem cell-like CD8^+^ T cells, or mediate mitochondrial changes that augment memory-like metabolic phenotypes. However, the molecular and metabolic mechanisms inducing the clear benefits of metabolic conditioning remain poorly understood.

Here, we show that culturing fully activated CD8^+^ effector T (T_E_) cells for 20 hours in media containing 1 mM glucose (here referred to as ↓[GLC]-conditioned T_E_ cells) (Figure 1A) increases mitochondrial electron transfer chain (ETC) complex III ROS (CIII ROS), shifting T_E_ cells to an oxidized cellular redox state. This redox shift leads to upregulated p38 signaling and increases transcription of effector molecules upon restimulation, augments *in vivo* persistence of adoptively transferred T_E_ cells and contributes to enhanced *in vivo* tumor clearance. In support of a role for beneficial redox changes, altering redox biology in *ex-vivo* expanded human tumor infiltrating lymphocytes (TILs) increased p38 phosphorylation and pro-inflammatory cytokine production. Collectively, our data suggest that the redox-p38 signaling axis provides a previously undiscovered mechanistic link between metabolic conditioning of T cells and anti-tumor T cell function and represents a targetable pathway that can be used to increase T cell-based anti-tumor immunotherapy efficacy.

**Figure. 1.**
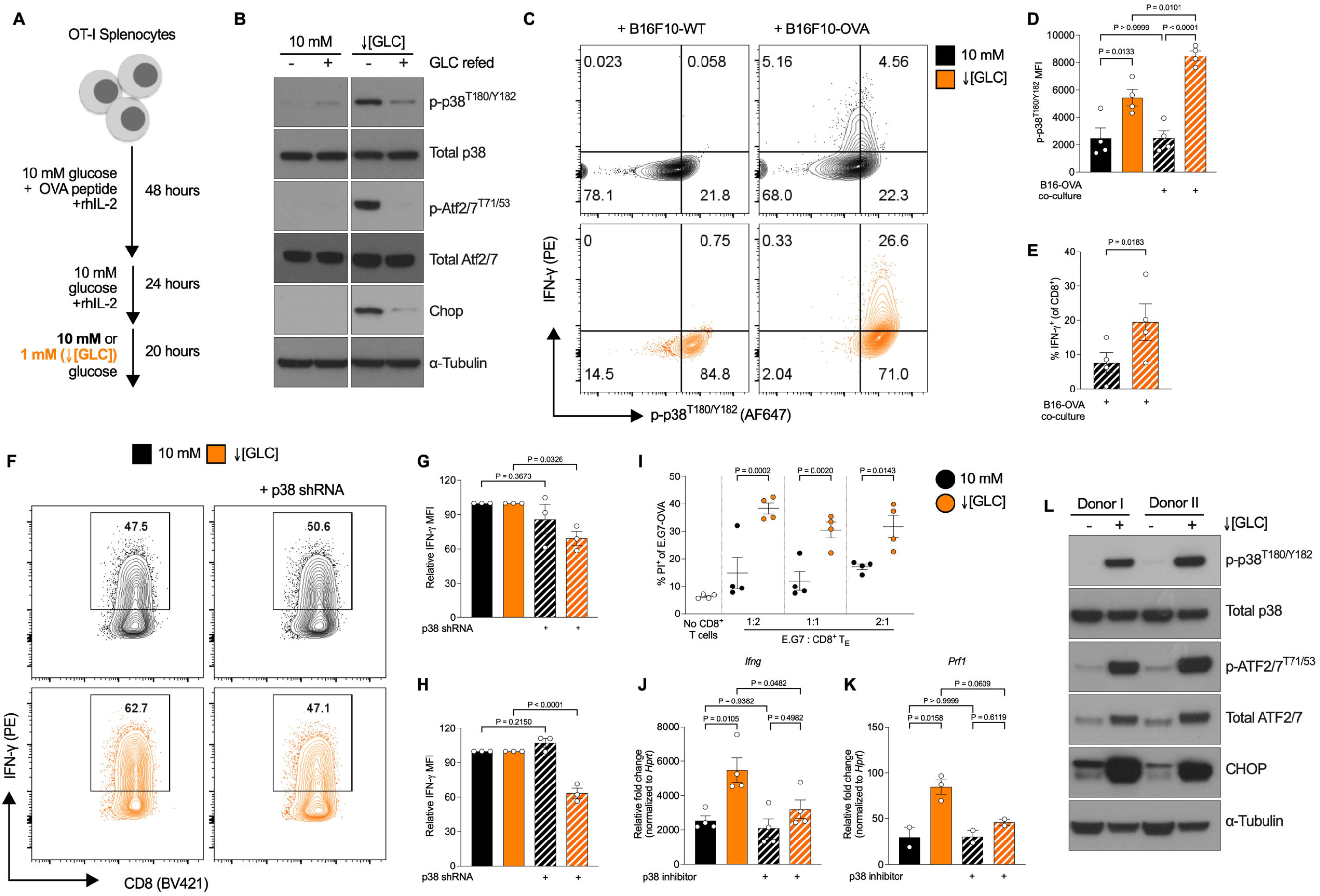
↓[GLC] conditioning increases p38-dependent CD8^+^ T cell function. (**A**) Schematic illustration of mouse CD8^+^ T cell activation with the indicated conditions; 10 mM glucose (control) and 1 mM glucose (↓[GLC]). (**B**) Representative immunoblot analysis of protein extracts isolated from control and ↓[GLC]-conditioned T_E_ cells with or without a 4-hour glucose refeed, probed for phosphorylated p38 at T180/Y182 (p-p38^T180/Y182^), total p38, phosphorylated ATF2/7 at T71/53 (p-Atf2/7^T71/53^), total Atf2/7, and Chop. α-Tubulin was used as a loading control. Data are representative of n > 6 independent experiments with 3 biological replicates each. (**C**) Representative flowcytometric contour plots of IFN-γ and p-p38^T180/Y182^ from OT-I T_E_ cell cocultures with either B16F10-WT or B16F10-OVA. n = 4 biologically independent samples, representative of >3 independent experiments (**D**) Quantification of p- p38^T180/Y182^ per cell (Mean fluorescence intensity; MFI) and (**E**) the frequency of IFN-γ^+^ cells are shown. (**F**) Representative flow cytometric contour plots of IFN-γ expression with or without p38 shRNA expression in 10 mM and ↓[GLC]-conditioned T_E_ cells after coculture with B16F10- OVA cell line. Quantification of (**G**) p-p38^T180/Y182^ MFI and (**H**) IFN-γ MFI. (**I**) Quantification of tumor cell killing by control and ↓[GLC]-conditioned T_E_ cells in cocultures with EG.7-OVA lymphoma cells after 24 hours. qRT-PCR analysis of (**J**) *Ifng* and (**K**) *Prf1* expression with or without a 4-hour αCD3/28 restimulation of control and ↓[GLC]-conditioned T_E_ cells. Representative of 2 independent experiments with 2 different p38 inhibitors. (**L**) Immunoblot analysis of protein expression for p-p38^T180/Y182^, total p38, p-ATF2/7^T71/53^, total ATF2/7, CHOP in fully activated control and ↓[GLC]-conditioned human CD8^+^ T_E_ cells. α-Tubulin was used as a loading control. n = 2 biological independent samples, representative of 3 independent experiments. Data are presented as mean ± SEM and *P* values are determined by two-tailed Student’s *t*-test or one-way ANOVA for more than 2 groups comparisons. See also Figure S1.

## RESULTS

### ↓[GLC] conditioning induces p38 kinase activation and enhances CD8^+^ T_E_ cell cytotoxicity

We previously showed that T_E_ cells cultured in ↓[GLC] condition *in vitro* drastically improved *in vivo* anti-tumor responses ^18^. Although we observed alteration in redox homeostasis and metabolic changes, the mechanism underlying the improved function remained elusive. ↓[GLC] induced a dramatic difference in cellular redox homeostasis as evidenced by a depletion of NADPH and accumulation of oxidized glutathione ^18^. To further our understanding of the underlying mechanism improving the *in vivo* tumor control, we next queried commonly activated pathways associated with glucose depletion and redox stress. Culturing mouse CD8^+^ T_E_ cells in ↓[GLC] induced a more oxidized cellular redox state and increased p38 phosphorylation after 20 hours (Figure 1B). Although a robust and reproducible increase in the phosphorylation of p38 was observed after 20 hours of culture, when glucose (10 mM) was added back to the cultures this was rapidly reversed (Figure 1B). Strikingly, and in agreement with our previous work, culture in ↓[GLC] for 20 hours showed no increase in phosphorylation of AMP kinase (AMPK), nor did we see an increase in phosphorylation of the stress integrating factor eukaryotic initiation factor 2a (eIF2a) (Figure S1A) suggesting that the depletion of glucose, glycolysis intermediates and a more oxidized cellular state did not induce canonical stress signaling through PERK, HRI or GCN2 which all converge on eIF2a ^22, 23^. We did not detect overall increases in protein oxidation/carbonylation (Figure S1B), suggesting that there is no complete loss of redox sufficiency leading to accumulated oxidized protein products. One potential compensatory pathway to sustain GSH biosynthesis in the absence of extracellular glucose is the enhanced uptake of glutamine and secretion of glutamate (Figures S1C, D), presumably for the transport of cystine by xCT/Slc7a11 ^24^. The increase of p38 signaling was accompanied by increased Atf2/7 phosphorylation (Figure 1B), and expression of Chop (encoded by the *Ddit3* gene), both of which were reversed by glucose refeeding (Figures 1B and S1E). Similar to Chop, Atf4 (encoded by *Atf4*) was induced in ↓[GLC]-conditioned T_E_ cells (Figures S1A, F). This potentially reflects a transcriptional adaptation to increase the expression of nutrient transporters such as the increase in CD98 ^18^, previously shown to be an Atf4 target ^25^, likely to compensate for the loss of glucose-derived carbons. Previous studies showed that p38 signaling is crucial for TCR- dependent activation of T cells ^26^. Thus, we next asked if the p38 signaling pathway was differentially impacted by restimulation of control or ↓[GLC]-conditioned T_E_ cells in complete media containing 10 mM glucose using either αCD3/28 or coculture with WT or ovalbumin (OVA)-expressing B16 melanoma cells to mimic tumor:T cell interactions. We observed a sustained increase in p38 signaling in ↓[GLC]-conditioned T_E_ cells (Figures 1C, D), accompanied by an increase in *Atf4* expression (Figure S1G), but the expression of *Ddit3* did not go up as a result of restimulation in the presence of glucose (Figure S1H). Together, these results suggest that ↓[GLC]-mediated p38 signaling and TCR-mediated restimulation in the presence of glucose, potentially induce different p38-dependent signaling pathways in ↓[GLC]-conditioned T_E_ cells.

To assess the functional impact of p38 signaling on cytokine production, we used genetic or pharmacological inhibition of p38 in the context of restimulation. ↓[GLC]-conditioned T_E_ cells produce more effector cytokines in tumor cocultures (Figure 1E, F), but this was reversed to levels in control T_E_ cells by shRNA-mediated knockdown of p38 (Figures 1F-G and S1I). Strikingly, the control T_E_ cells did not show a reduction, but rather slightly augmented interferon-gamma (IFN-γ) production during αCD3/28-mediated restimulation (Figure 1H), corroborating the previous observation that inhibition of p38 in CD8^+^ T cells can augment their anti-tumor function in the context of glucose sufficiency ^27^. In support of increased T_E_ function, ↓[GLC]-conditioned T_E_ cells also significantly enhanced tumor cell killing of OVA-expressing E.G7 lymphoma cells *in vitro* at various T_E_ cell to tumor cell ratios (Figure 1I). Increased p38 signaling was associated with increased transcription of the effector molecules perforin (*Prf1*) and interferon gamma (*Ifng*), suggesting that the increase in effector function of these cells could be at least in part transcriptionally mediated (Figures 1J, K).

To extend these observations from mouse to human CD8^+^ T cells, we next isolated naïve CD8^+^ T cells from blood donated by healthy individuals, followed by activation using αCD2/3/28 tetramers for 5 days in proprietary ImmunoCult media (STEMCELL Technologies) containing 200 U/ml rhIL-2 to generate rapidly proliferating CD8^+^ T_E_ cells. Cells were then split into serum-free, 10 or 1 mM glucose containing custom ImmunoCult media overnight. Cells exposed to ↓[GLC] (indicated by complete glucose depletion in the cultures (Figure S1J)) significantly upregulated p38 phosphorylation and downstream activation of ATF2/7 (Figure 1L), suggesting that the glucose depletion-induced p38 signaling pathway is conserved between mouse and human CD8^+^ T_E_ cells.

### Mitochondrial ETC complex III ROS induces p38 activation in **↓**[GLC]-conditioned T_E_ cells

To uncover the mechanism leading to increased p38 activity in ↓[GLC]-conditioned T_E_ cells, we hypothesized that the altered redox balance was upstream of p38 signaling. Thus, we treated T_E_ cells in ↓[GLC] cultures with various ROS scavengers and redox modulators. Addition of N-acetyl-L-cysteine (NAC), a reduced glutathione precursor, did not affect p38 phosphorylation (Figure 2A); the same was true for cell-permeable methylated glutathione (GSH-MEE) (Figure S2A). On the other hand, addition of the pentose phosphate pathways (PPP) inhibitor, 6-AN, markedly increased p38 phosphorylation (Figure S2B). 6-AN-mediated PPP inhibition, associated with GSSG accumulation and NADPH depletion, was previously utilized in OT-I T cells to enhance *in vivo* anti-tumor T cell function ^19^. Taken together, these results suggested to us that decreased reducing equivalents in the form of NADPH and the resulting oxidized redox status are upstream of the increased p38 activity. To assess mechanistic drivers of the altered T_E_ cell redox status, we examined the contribution of mitochondrial ETC complex III (CIII), a source of signaling ROS important for full T cell activation ^28^. Although not significant across the whole population^18^ (Figure S2C), MitoSOX staining which detects mitochondrial superoxide, reproducibly showed an increase in the mitoSOX^high^ population in the ↓[GLC]-conditioned T_E_ cells (Figure 2B). This was not the case for CellROX staining, which measures cellular ROS (Figure S2D). Addition of S3QEL-2, a specific inhibitor of superoxide production from mitochondrial ETC CIII that does not interfere with ETC function ^29^, reduced the mitoSOX^high^ population (Figure 2B) and down-regulated p38 phosphorylation in ↓[GLC]-conditioned T_E_ cells both before and after TCR-mediated restimulation (Figure 2C). Since p38 signaling augmented cytokine production in ↓[GLC]-conditioned T_E_ cells (Figure 1), we next asked if blunting mitochondrial ETC CIII ROS would affect cytokine production. As shown above, ↓[GLC]- conditioned T_E_ cells produce significantly more IFN-γ in cocultures with B16-OVA tumor cells, a phenotype that was abrogated by S3QEL-2 treatment during the ↓[GLC]-conditioning culture (Figures 2D,E). Similarly, αCD3/28 restimulation enhanced IFN-γ production in ↓[GLC]- conditioned T_E_ cells, which was also blunted by inhibition of mitochondrial ETC CIII ROS (Figures 2F,G). As a further link to an altered cellular redox state, we observed that ↓[GLC]- conditioned T_E_ cells displayed a further oxidized cytosol after 4 hours of restimulation with αCD3/28 (Figure 2H), which corresponds to the increased CIII ROS-mediated p38 signaling (Figure 2C) and cytokine production (Figures 2D-G). Taken together, these results suggest that mitochondrial ETC CIII ROS contributes to 1) enhanced p38 signaling, 2) enhanced cytokine production, and 3) a more oxidized subcellular redox state in ↓[GLC]-conditioned T_E_ cells.

**Figure. 2.**
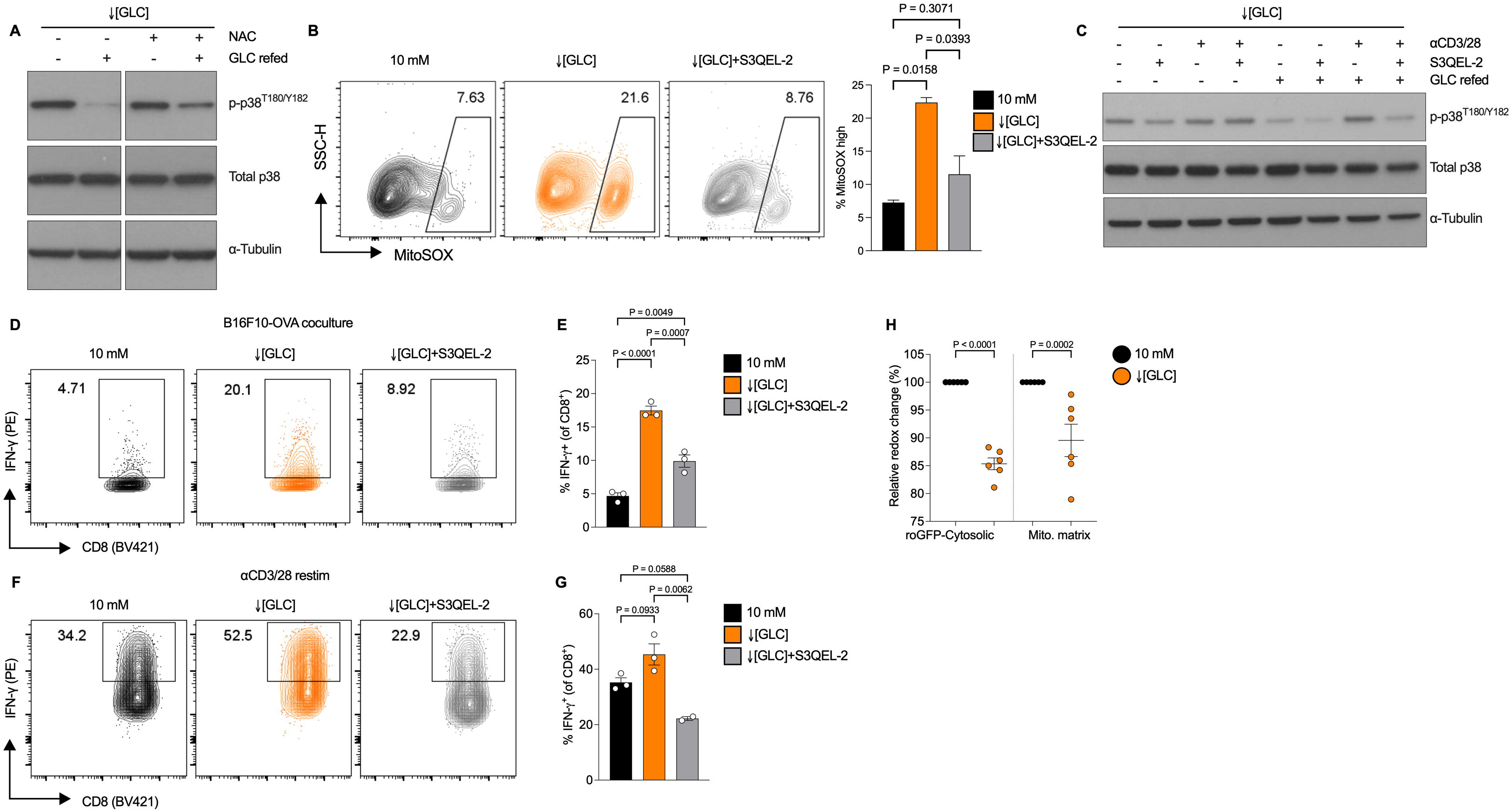
p38 signaling and cytokine production in↓[GLC]-conditioned T_E_ cells is increased by mitochondrial complex III ROS. (**A**) Representative immunoblot analysis of p- p38^T180/Y182^ and total p38 in control and ↓[GLC]-conditioned T_E_ cells with or without N-acetyl- L-cysteine (NAC, 10 mM) during the 20-hour culture, followed by a 4-hour glucose refeed. Representative of 3 biological independent samples. (**B**) Quantification of the frequency of the mitoSOX^high^ population and representative flowcytometric contour plots of mitoSOX staining (**C**) Representative immunoblot analysis of p-p38^T180/Y182^ and total p38 in control and DMSO or S3QEL-2 treated ↓[GLC]-conditioned T_E_ cells with or without restimulation for 4 hours with αCD3/28. n = 2 biological independent samples. (**D**) Flow cytometric analysis of IFN-γ expression after coculture of B16F10-OVA and control or↓[GLC]-conditioned T_E_ cells treated with DMSO or S3QEL-2. (**E**) Quantification of the frequency of IFN- ^+^ (of CD8^+^ T cells) from d). (**F**) Flow cytometric analysis of IFN-γ expression after αCD3/28 restimulation of CD8^+^ T_E_ cells as in d). (**G**) Quantification of the frequency of IFN- ^+^ (of CD8^+^ T cells) from f). (**H**) roGFP-based analysis of relative cytosolic redox state in control and ↓[GLC]-conditioned T_E_ cells after 4 hours of restimulation with αCD3/28 beads. n = 6 biologically independent samples. Data are presented as mean ± SEM and *P* values are determined by two-tailed Student’s *t*-test or one-way ANOVA for > 2 groups comparisons. See also Figure S2.

### Altered redox homeostasis in **↓**[GLC]-conditioned T_E_ cells enhances T cell function

We observed that S3QEL-2 treatment during ↓[GLC]-exposure partially reverted the subcellular redox status as measured by cytosolic (Figure 3A) and mitochondrial roGFP (Figure 3B) towards a more homeostatic level seen in control T_E_ cells. Next, we asked if ETC CIII ROS-mediated p38 signaling (Figure 2C) was directly linked to the altered redox-couple state. We treated CD8^+^ T_E_ cells with S3QEL-2 during ↓[GLC] conditioning and observed that dampening ETC CIII ROS significantly increased the cellular NADPH/NADP^+^ (Figure 3C) and GSH/GSSG ratios (Figure 3D) in↓[GLC]-conditioned T_E_ cells, shifting them to a less oxidized phenotype. We observed that the differences in GSH/GSSG ratios of ↓[GLC]-conditioned T_E_ cells and the reversal by S3QEL-2 treatment was driven by changes in the levels of the oxidized form of glutathione (GSSG) abundance (Figure 3E) rather than the pools of reduced GSH (Figure 3F), further linking this phenotype to levels of NADPH that are crucial for the reduction of oxidized GSSG and crucial for reductive biosynthetic metabolic pathways. To test if reduced levels of NADPH induced by ↓[GLC] conditioning (Figure 3C) could be directly enhancing cytokine production, we retrovirally expressed an exogenous NADPH oxidase, TPNOX, which consumes NADPH and generates H_2_O but not ROS ^30^. Expression of TPNOX increased p38 signaling in the presence of glucose (Figure S3A) and markedly increased IFN-γ production in restimulated T_E_ cells in 10mM glucose cultures (Figures 3H,I), suggesting a direct link between NADPH homeostasis and T_E_ cell function.

**Figure. 3.**
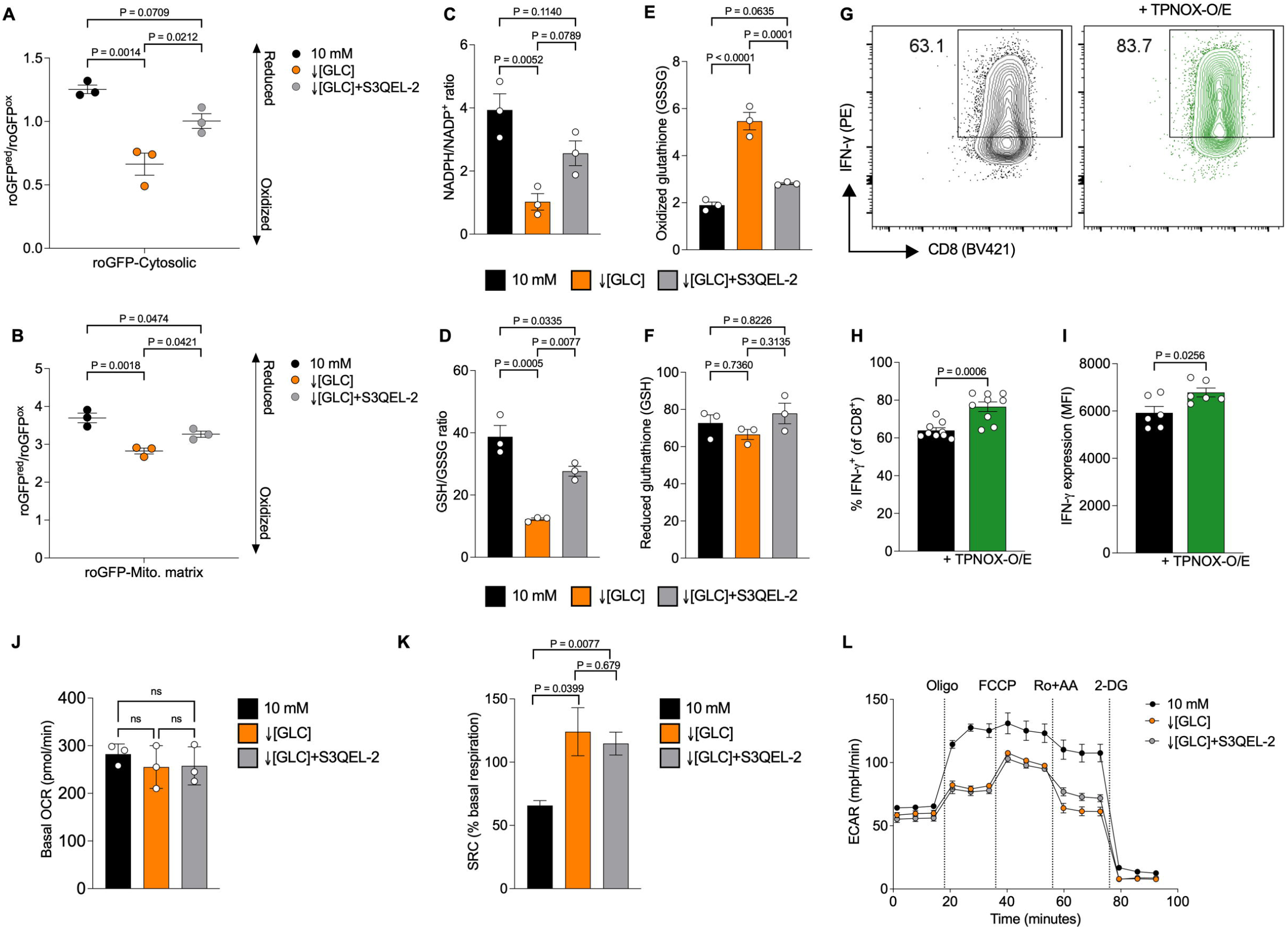
Altered redox homeostasis enhances T_E_ cell function. roGFP-based analysis of relative redox state in the (**A**) cytosol and (**B**) mitochondrial matrix of control and DMSO or S3QEL-2 treated ↓[GLC]-conditioned T_E_ cells (**C**) NADPH/NADP+ ratio (**D**) GSH/GSSG ratio (**E**) oxidized GSSG pool and (**F**) reduced GSH pool of control and DMSO or S3QEL-2 treated ↓[GLC]-conditioned T_E_ cells (**G**) Representative flowcytometric contour plots of IFN-γ expression after αCD3/28 restimulation of CD8^+^ T_E_ cells with or without TPNOX overexpression (O/E) grown in 10 mM glucose. (**H**) Quantification of the frequency of IFN-γ ^+^ cells and (**I**) IFN-γ MFI from cells in g). n = 6 biological independent samples are shown. Mitochondrial stress test using a Seahorse extracellular flux analyzer showing (**J**) mitochondrial respiration represented by oxygen consumption rate (OCR) (**K**) Quantification of spare respiratory capacity (SRC) as a percentage of the basal respiration and (**L**) glycolytic activity represented by extracellular acidification rate (ECAR) in control and DMSO or S3QEL-2 treated ↓[GLC]-conditioned T_E_ cells. Data are presented as mean ± SEM and *P* values are determined by two-tailed Student’s *t*-test or one-way ANOVA followed by multiple comparisons for > 2 groups comparisons. See also Figure S3.

Given the observed role for ETC CIII ROS in the induction of redox alterations and p38 signaling, we assayed the relative mitochondrial activity of control and ↓[GLC]-conditioned T_E_ cells. As we previously observed, ↓[GLC] induces mitochondrial elongation and spare respiratory capacity (SRC) ^31^, which was suggested to contribute to T cell persistence *in vivo* ^32^ and anti-tumor and memory T cell function ^31, 33^. Inhibiting mitochondrial ETC CIII ROS in ↓[GLC]-conditioned T_E_ cells did not affect mitochondrial respiration (Figure 3J) and maintained the induction of mitochondrial SRC (Figure 3K), nor did S3QEL-2 treatment affect glycolysis as assessed by Seahorse extracellular flux analysis graphs that are essentially identical (Figure 3L). We did observe that glucose remained detectable in S3QEL-2-treated ↓[GLC]-conditioned T_E_ cell culture supernatants (Figure S3B despite similar ECAR traces), while lactate secretion was comparable to DMSO treated ↓[GLC]-conditioned T_E_ cells (Figure S3C). Together these data suggest that ↓[GLC]-induced ETC CIII ROS or TPNOX expression contribute to changes in cellular redox and p38 signaling, augmenting T_E_ cell function, without alterations in mitochondrial respiration and mitochondrial SRC or cell intrinsic glycolysis.

### Inhibition of mitochondrial ETC complex III ROS blunts the enhanced anti-tumor activity of **↓**[GLC]-conditioned T_E_ cells

Finally, we asked if ETC CIII ROS-mediated redox changes that drive p38 activity contributed to the improved tumor clearance *in vivo* we previously observed with ↓[GLC]-conditioned T_E_ cells ^18^. Immunocompetent female WT C57BL/6J recipient mice (CD45.1^+^) were implanted with 1 × 10^6^ OVA-expressing EL4 (E.G7-OVA) lymphoma cells subcutaneously. Eight days after tumor implantation, recipient mice were randomized and 3 ×10^6^ OVA-specific control CD8^+^ T_E_ cells, DMSO or S3QEL-2 treated ↓[GLC]-conditioned T_E_ cells (CD45.2^+^, Figure 4A) were washed and resuspended in drug-, serum-, and glucose-free media and injected intravenously. All (7/7) mice that received no OVA-specific T cells failed to control tumor growth and had to be sacrificed about 18 days after tumor implantation (Figure 4B open circles). We observed that adoptive transfer of ↓[GLC]-conditioned T_E_ cells (Figure 4B Orange lines) cleared tumors significantly better than control CD8^+^ T_E_ cells (Figure 4B Black lines). Strikingly, blunting ETC CIII ROS with S3QEL-2 during *in vitro* ↓[GLC] conditioning significantly impacted the ability of the cells to control tumor growth (Figure 4B Grey lines), despite the drug being washed out before injection. Key features of therapeutic T cell products are *in vivo* persistence, tumor infiltration, resistance to T cell exhaustion and sustained *in vivo* anti-tumor effector function ^34^. ↓[GLC]-conditioned T_E_ cells persisted longer in circulation than control T cells both 6 and 15 days after the adoptive transfer (Figure 4C), but strikingly SQ3EL-2 treatment under ↓[GLC] condition in CD8^+^ T_E_ cells before injection significantly reduced the numbers of circulating CD8^+^ CD45.2^+^ donor cells (Figure 4C). We also analyzed donor cell percentages in the spleen and tumor draining lymph nodes (tdLN), and found that as in the blood, ↓[GLC] conditioning augmented persistence in the spleen, but surprisingly not the tdLN of these mice (Figures S4A, B). The augmented persistence in the spleen was lost by blocking ETC CIII ROS (Figure S4A). In support of a sustained functional link between ↓[GLC] culture and p38 phosphorylation induced *in vitro* in ↓[GLC]-conditioned T_E_ cells, p38 phosphorylation in donor CD8^+^ T cells isolated from blood was still increased by restimulation 15 days after infusion (Figure 4D). Of note, restimulation-mediated p38 signaling was no longer dampened by *in vitro* S3QEL-2 treatment during ↓[GLC] culture 15 days earlier (Figure 4D). Similarly, after restimulation, IFN-γ expression was also increased in the ↓[GLC]-conditioned donor CD8^+^ T_E_ in the blood compared to control donor CD8^+^ T_E_ cells (Figure 4E), but like p38 signaling, IFN-γ production was no longer blocked by the *in vitro* S3QEL-2 treatment during ↓[GLC] conditioning. Similar to the smaller tumor diameter in mice receiving ↓[GLC]-condition T_E_ cells (Figure 4B), we observed a significant reduction in tumor weight in both ↓[GLC] groups when compared to control CD8^+^ donor cells (Figure 4F). After digestion of the tumors, we stained for donor cells and found that ↓[GLC] CD8^+^ donor T cells infiltrated the tumors significantly better (Figure 4G). However, S3QEL-2 treatment before infusion blunted increased infiltration, since the number of S3QEL-2-treated -↓[GLC] conditioned T_E_ cells was no longer significantly higher than those of the control CD8^+^ T cells (Figure 4G). Of note, there is no significant difference in tumor infiltration between the two ↓[GLC]-conditioned T_E_ cell groups (Figure 4G), despite the S3QEL- 2-treated CD8^+^ T_E_ cells during *in vitro* ↓[GLC] conditioning having a significant lower number of donor CD8^+^ T cells in circulation (Figure 4C). This suggests that the enhanced tumor clearance of ↓[GLC]-conditioned T_E_ cells could be either due to better persistence in circulation or linked to enhanced function inside the tumors that is supported by ETC CIII ROS. To test these possibilities, we next restimulated tumor-infiltrating CD8^+^ T cells with PMA/ionomycin and measured the expression of p-p38 and IFN-γ. We observed that the ↓[GLC]-conditioned donor CD8^+^ T_E_ cells expressed significantly higher p-p38 (Figure 4H) and a trend towards increased IFN-γ (Figure 4I) after restimulation when compared to control donor CD8^+^ T cells. *In vitro* inhibition of ETC CIII ROS led to a loss of the ↓[GLC]-induced increase in p-p38 (Figure 4H) and a trend towards lower IFN-γ in tumor-infiltrating donor CD8^+^ T cells (Figure 4I).

**Figure. 4.**
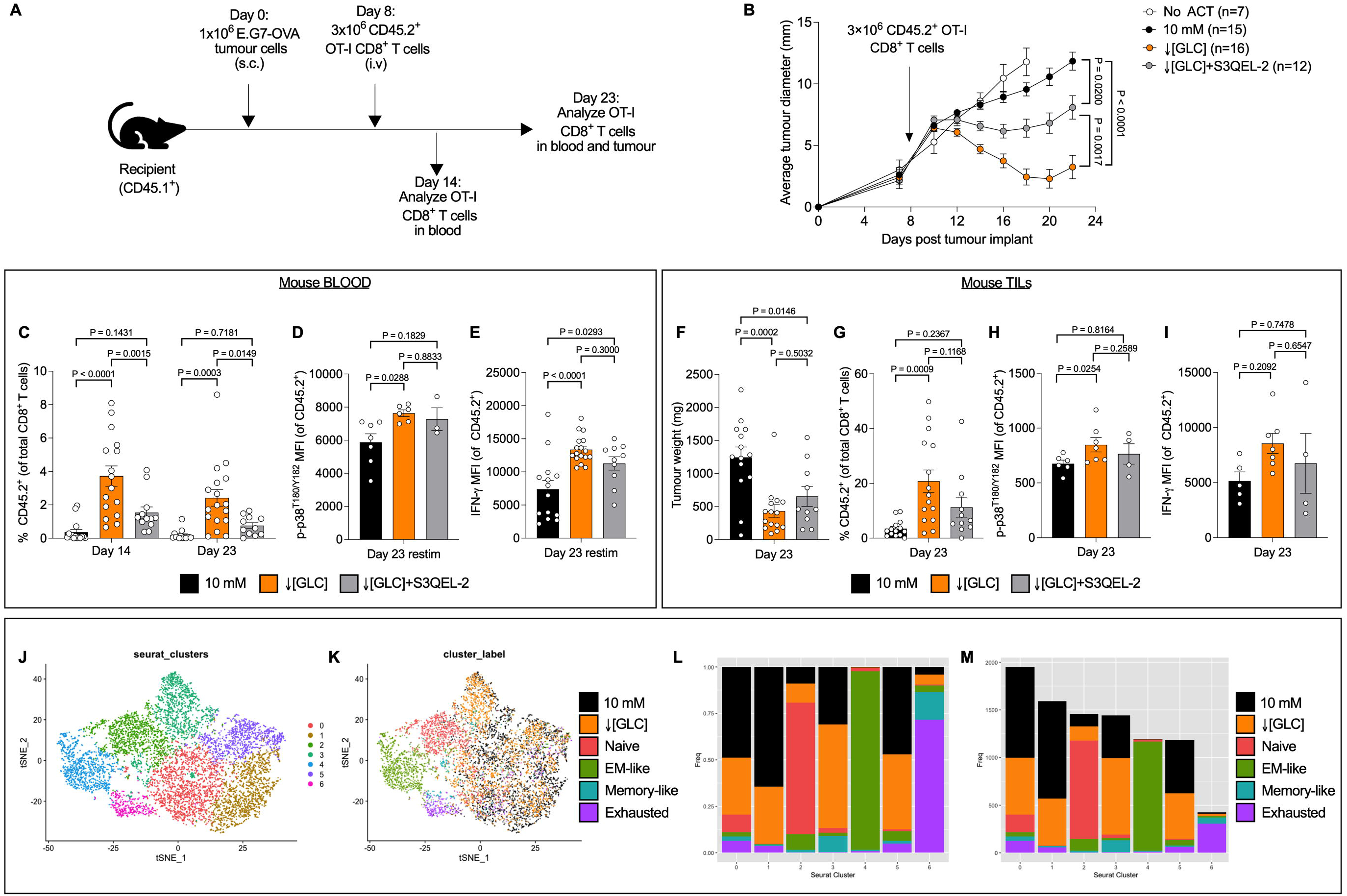
Mitochondrial complex III ROS is a key contributor to the enhanced anti-tumor function of ↓[GLC]-conditioned T cells *in vivo*. (**A**) OT-I splenocytes from 8-12-week-old female CD45.2 mice were activated with SIINFEKL peptide and expanded for 72 hours in media containing 10 mM glucose and IL-2, followed by exposure to 10 mM or 1 mM glucose (↓[GLC]) with or without S3QEL-2 for 20 hours. 1 × 10^6^ E.G7-OVA lymphoma cells were injected in the flank of immunocompetent female CD45.1 mice and allowed to grow for 8 days. 3 × 10^6^ OT-I T_E_ cells were intravenously injected per tumor-bearing mouse. (**B**) Average diameter of E.G7- OVA lymphoma in mice injected receiving no TE cells (open circle; n = 7), control T_E_ (black circle; n = 15), ↓[GLC]-conditioned T_E_ (orange circle; n = 16), or S3QEL-2 pre-treated ↓[GLC]- conditioned T_E_ (grey circle; n = 12). *P* values are determined by two-way ANOVA with multiple comparisons. (**C**) Blood was collected 6 and 15 days post T_E_ cell transfer and the frequency of CD45.2^+^ donor CD8^+^ T cells was measured by flowcytometry. (**D**) Expression of p-p38^T180/Y182^ per cell (MFI), and (**E**) IFN-γ MFI of CD45.2^+^ donor CD8^+^ T cells derived from the blood were measured after *ex vivo* restimulation with PMA/ionomycin for 4 hours. (**F**) Quantification of tumor weight (mg) at the experimental endpoint (Day 23). (**G**) Frequencies of CD45.2^+^ donor CD8^+^ T cells as a percentage of total CD8^+^ cells in tumors were assessed by flow cytometry. (**H**) Expression of p-p38^T180/Y182^ per cell (MFI), and (**I**) IFN-γ MFI of CD45.2^+^ donor CD8^+^ tumor infiltrating T cells isolated from the tumors were measured after *ex vivo* restimulation with PMA/ionomycin for 4 hours. (**J, K**) tSNE plots were generated using the first 10 principal components, of the combined datasets of control and ↓[GLC]-conditioned T_E_ cells and melanoma infiltrating CD8^+^ T cells. For clustering, shared nearest neighbor modularity optimization clustering algorithm from Seurat v3 was used with resolution parameter = 0.5. A total of 7 clusters were obtained. Left panel showing the 7 clusters, right panel showing cell identity labels on the same tSNE plot, indicating cluster 3 as enriched in ↓[GLC]-conditioned T_E_ cells. Representation of the frequency (**L**) and count (**M**) of cell identities per cluster. Cluster 3 shows enrichment of both ↓[GLC]-conditioned T_E_ cells and Memory Like tumor infiltrating T cells. Data in panels C-I are presented as mean ± SEM and *P* values are determined by one-way ANOVA, followed by multiple comparisons. See also Figure S4.

### Distinct immunological phenotypes in of ↓[GLC]-conditioned TE cells

Breakthrough studies have shown that transcriptional and epigenetic signatures that correlate with better *in vivo* persistence and anti-tumor T cell function can be identified, and have led to the discovery of memory-like and stem-cell like CD8^+^ T cell subsets with enhanced anti-tumor capacity ^35, 36^. Similarly, selection of cells with low mitochondrial membrane potential^17^ or low glucose metabolism ^16^, inducing changes in mitochondrial mass by genetic or pharmacologic strategies ^15, 37^, or the alterations of mitochondrial morphology and ultrastructure^31, 33^, are all associated with memory-like CD8^+^ T cell subsets and have been shown to augment anti-tumor function in preclinical models. We did not see phenotypic evidence of memory differentiation in the transferred ↓[GLC]-conditioned T_E_ cells in the tumor draining lymph node (tdLN) or tumor, where CD62L expression remained unchanged from the control T_E_ cells (Figures S4C, D). Given the obvious link of memory-like CD8^+^ T cell subsets with beneficial outcomes, we queried our previously published single cell RNA sequencing data from *in vitro* control and ↓[GLC]- conditioned T_E_ cells ^18^ for overlap with stem-cell like CD8^+^ T cell transcriptomes observed in B16 melanoma models ^38^. Surprisingly, despite obvious transcriptomic heterogeneity across clusters (Figures 4J-M), we found that of the *in vitro* cultured ↓[GLC]-conditioned T_E_ cells, a subset clustered closely with memory-like TILs isolated from B16 melanoma tumors (Cluster 3, Figures 4J-M). Strikingly, cluster 3 is the only cluster that shows lower enrichment of the control T_E_ cells (Figure 4L, M). This finding suggests that beyond the lower expression of memory-like surface markers like CD62L (Figure S4E) and the maintenance of effector T cell markers such as CD25 and CD98^18^, there are transcriptional modules that could be mediated by the metabolic conditioning of ↓[GLC]-conditioned T_E_ cells such as targets of the p38 pathway in addition to the cytokines we showed to be induced in this study.

### Induction of redox cycling in human CD8^+^ TILs increases IFN-**γ** production

Since *in vitro* activated human CD8^+^ T_E_ cells from healthy individuals induced enhanced p38 signaling in ↓[GLC] cultures (Figure 1L), we next addressed if the effector function of tumor infiltrating CD8^+^ T cells isolated from human keratinocyte-derived skin cancer (KDSC) punch biopsies could be augmented by the induction of redox imbalance and/or p38 signaling. In KDSC, multiple immunosuppressive conditions were associated with increased risk, indicating a strong association with the immune system ^39^, and immune checkpoint therapy is rapidly evolving as a promising therapeutic strategy ^40^. Histologically, a marked immune cell infiltrate, including T cells, represents a hallmark of KDSC ^41^. Consistently, we observed increasing numbers of CD3^+^ T cells towards the tumor center (Figure S5A). Critically, CD8^+^ TILs lost mitochondrial content in the tumor as assessed by a loss of MitoTracker green staining compared to CD8^+^ T cells isolated from adjacent skin from the same patient *ex vivo* (Figure 5A). CD8^+^ TILs from human KDSC show reduced p-p38^T180^ expression compared to CD8^+^ T cells from adjacent skin *ex vivo* (Figure 5B), perhaps associated with reduced mitochondrial ROS-mediated redox signaling. To test this, total CD3^+^ T cells were isolated from tumor biopsies, and expanded through restimulation with αCD2/3/28 tetramers for 5 days (Figure 5C). To assess effector function, TILs were restimulated with αCD2/3/28 tetramers for 5 hours *ex vivo* which showed limited production of the effector cytokine IFN-γ from the CD8^+^ TIL population (Figure 5D left panel). To force a redox imbalance in these cells, in an effort to induce beneficial p38 activity, we pre-treated the TIL cultures with the redox cycling compound 2,3-dimethoxy-1,4- naphthoquinone (DMNQ) ^42^ before restimulation. DMNQ induces an increase in GSSG, which leads to reduced NADPH pools due to glutathione reductase activity to preserve GSH pools, similar to what we observed in ↓[GLC]-conditioned T_E_ cells (Figures 2 and 3). Strikingly, DMNQ treatment induced p-p38 expression in the expanded KDSC TIL cultures isolated from all patients (Figure 5E), suggesting that the redox mediated p38 pathway is conserved in human CD8^+^ TILs. In agreement with our findings in mouse CD8^+^ T_E_ cells by ↓[GLC]-conditioning or TPNOX-mediated depletion of NAPDH, the induction of redox-mediated p38 signaling augmented IFN-γ production in CD8^+^ TILs (Figure 5F), where the p-p38^high^ population was enriched with cells that recovered the ability to produce IFN-γ (Figure 5D right panel). Similar increased effector function was observed in fully activated mouse OT-I CD8^+^ T cells grown in 10mM glucose, where DMNQ treatment before either coculture with B16F10-OVA melanoma cells (Figure S5B) or αCD3/28 restimulation (Figure S5C) markedly increased IFN-γ production. Taken together these findings suggest that metabolic conditioning by either ↓[GLC] culture leading to redox imbalance-mediated p38 activation, or direct induction of redox imbalance leading to p38 activation in TILs, could present new possibilities to improve CD8^+^ T cell immunotherapy of cancer.

**Figure 5.**
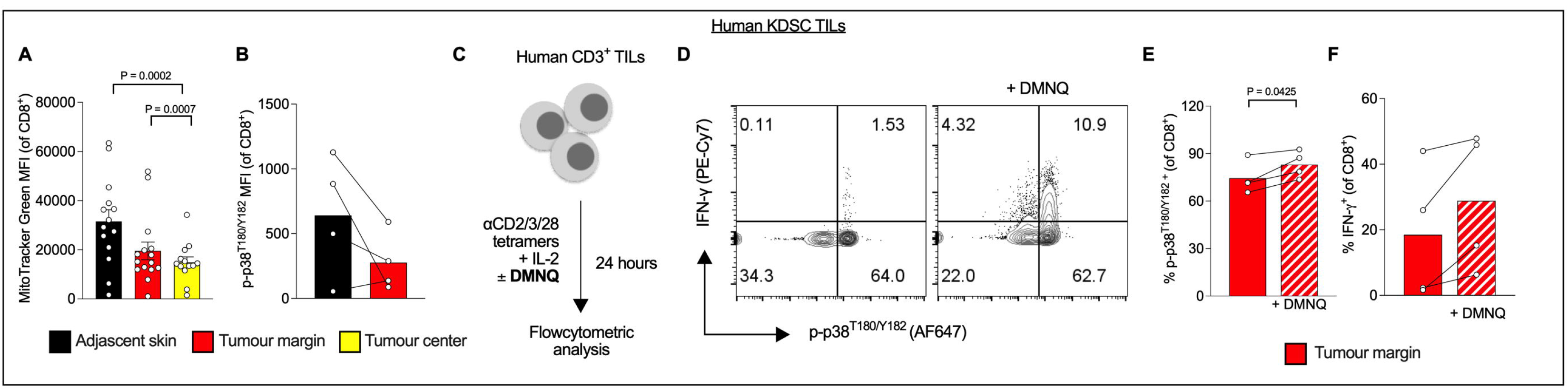
Induction of redox cycling in human CD8+ TILs increases IFN-γ production. (**A**) CD8^+^ T cells isolated from punch biopsies of adjacent skin, tumor-margin or -center were assessed for mitochondrial content using MitoTracker Green n = 15 biological independent samples. (**B**) analysis of p38 phosphorylation in CD8^+^ T cells isolated from matched adjacent and tumor margin skin, (**C**) CD3^+^ tumor infiltrating T cells were isolated from KDSC tumors, expanded by stimulation with αCD2/3/28 tetramers, followed by restimulation with αCD2/3/28 tetramers in the absence or presence of DMNQ (1 μM) for 24 hours. (**D**) Representative flowcytometry contour plot showing p-p38^T180/Y182^ and IFN-γ in restimulated TILs treated with or without DMNQ. (**E**) Quantification of p-p38^T180/Y182^ and (**F**) IFN-γ expression in restimulated CD8^+^ TILs pre-treated with or without DMNQ. Data in panels A-F are presented as mean ± SEM and statistical significance is determined by paired *t*-test. See also Figure S5.

## DISCUSSION

Previous studies have shown that glucose metabolism ^18, 43–47^, mitochondrial function ^37, 48, 49^, dynamics, morphology and SRC ^31–33^, ETC CIII ROS ^28^, NADPH generation ^19, 50^, GSH synthesis ^51^, and p38 signaling ^26^ all contribute to T cell function. However, the intersection of glucose availability, mitochondrial SRC and ROS, NADPH and GSH homeostasis, the p38 signaling pathway *and more importantly*, their combined effects on anti-tumor T_E_ cell function remained incompletely understood. Given the vast changes of T cell function described in the studies listed above, a better understanding of the intersection of these phenotypes is crucial for the development of metabolic strategies that improve T cell function in the TME. Given the pleotropic role of p38-signaling in mammalian cells, it remains unclear what causes the opposing effects of p38 deletion in control and ↓[GLC]-conditioned T_E_ cells. p38 acts as a pro-inflammatory factor downstream of the TCR in ↓[GLC]-conditioned T_E_ cells, but can cause apoptosis when hyperactivated ^52^, perhaps explaining the beneficial effects of p38 deletion in glucose sufficient T_E_ cells also described in a previous study ^27^. Similar to p38, ROS can be beneficial and detrimental for T cell function. During T cell activation ETC CIII ROS is produced but T cells are equipped with a ROS buffering system via Gclc-mediated GSH synthesis to avoid ROS toxicity ^28, 50^. Our data suggest that the NADPH/NADP+ ratio is regulated by mitochondrial ETC CIII ROS in T_E_ cells as a result of ↓[GLC] conditioning. A previous study showed that PPP-mediated NADPH production during initial activation was crucial for cytokine production in T cells, a phenotype that could not be rescued by N-acetyl-L-cysteine (NAC) supplementation. Surprisingly, we found that NAC and GSH-MEE did not inhibit p38 phosphorylation in ↓[GLC]-conditioned T_E_ cells. Taken together this suggests that the reduced glutathione pools and/or blocking ROS itself are likely not affecting p38 phosphorylation directly in fully activated ↓[GLC] conditioned T_E_ cells. Although this data together with TPNOX-mediated NADPH depletion strongly suggests that lowering the NADPH levels in fully activated CD8^+^ T_E_ cells improves anti-tumor function, the exact molecular mechanism linking this to p38 signaling will be the subject of future studies in the laboratory. Metabolic optimization by gene editing strategies employing CRISPR-Cas9 to engineer better metabolically equipped T cells is being explored and has shown efficacy in increasing anti-tumor T_E_ cell function in various preclinical cancer models ^27, 53, 54^, but one drawback with such strategies is the potential loss of metabolic flexibility due to a genetic alteration, which could impact the ability of a T cell to adapt to diverse environments encountered from the circulation and secondary lymphoid organs to the TME. Strategies involving *in vitro* depletion of key T cell bioenergetic substrates such as glucose ^18^ and glutamine ^55^ or the increase in lactate ^20^ or potassium ^21^ to mimic the nutrient-depleted, waste-product-dense TME have improved T cell function and tumor control without genetically locking the T cells in a given metabolic state. However, the mechanistic understanding of what makes a perfect tumor-killing T cell with sustained *in vivo* persistence remains elusive.

Although ↓[GLC]-conditioning induced mitochondrial elongation and the induction of mitochondrial SRC^31^, it remains unclear whether ↓[GLC]-conditioned T_E_ cells differentiated into other CD8^+^ T cell subtypes upon transfer in tumor-bearing mice. Surprisingly we found that SRC in the absence of ETC CIII ROS was not able to sustain the increased *in vivo* persistence of ↓[GLC]-conditioned T_E_ cells. A strongly debated phenotypic measure of mitochondrial efficiency – SRC - remains incompletely understood in the context of *in vivo* T cell survival and anti-tumor function. Previous studies showed a correlation with SRC and *in vivo* persistence and memory T cell development^31–33^, yet other studies showed that effector memory T cells without mitochondrial SRC could mediate long-term protection in mouse models of viral infection^56^. The work presented in this study supports both the observation that SRC can correlate with *in vivo* persistence and function, but also that SRC alone does not appear to be sufficient in the absence of ETC CIII ROS. Perhaps previous studies utilizing T cell-specific deletion of von Hippel-Lindau (VHL), leading to stabilization of HIF-1α - a key transcription factor for hypoxia-induced glycolysis^56^ – which can also be stabilized by ETC CIII ROS^57^, could be enhancing similar phenotypes we observe in our ↓[GLC]-conditioned T_E_ cells independent of SRC itself. However, this could also suggest that ETC CIII ROS is induced in T_E_ cells with increased SRC, thereby increasing *in vivo* longevity, but this will require further investigation.

The bimodal distribution of mitoSOX staining in our *in vitro* cultures (Figure 2b) suggests that there is a subset of T_E_ cells that have increased ETC CIII ROS accumulation in both control and ↓[GLC]-conditioned T_E_ cells. However, the functional increase in either tumor cocultures or by restimulation does not align with the proportional increase in this mitoSOX^high^ population, suggesting that merely selecting for mitoSOX^high^ cells might not be sufficient to select T_E_ cells with superior anti-tumour function. Given the known issues with ROS detecting reagents, and off target effects of anti-oxidants^58^, we chose to further assess the role of redox changes using exogenous TPNOX expression, which does not increase cellular ROS, but consumes the redox co-factor NADPH. This further corroborated that there is heterogeneity in the cultures, since not 100% of the TPNOX-expressing T_E_ cells increased pro-inflammatory cytokine production. In an effort to further resolve the heterogeneity, we chose to re-analyse previously published scRNA sequencing data, showing that there is transcriptional heterogeneity in the *in vitro* generated T_E_ cells. A subpopulation of ↓[GLC]-conditioned T_E_ cells appears to be transcriptionally similar to the previously identified memory-like CD8^+^ T cells that are associated with superior anti-tumor function. Future studies focussing on subpopulations within the *in vitro* metabolically conditioned cultures might identify key metabolic, epigenetic and/or transcriptional changes that pre-dispose T_E_ cells to differentiate into therapeutically relevant beneficial T cell subsets.

A by-product of oxidative metabolism in mitochondria, ROS can lead to damage of RNA, DNA, and proteins, potentially leading to cell death^59^. ROS are implicated as both detrimental and beneficial molecules in both tumour and T cells^60, 61^. These conflicting data suggest the impact of ROS is context-dependent, and affects both tumour cells and anti-tumour T cell responses. The development of therapeutic strategies to limit ROS in ROS-dependent tumours, could lead to a reduction of T cell function, but equally an increase in ROS to induce cancer cell death, could lead to T cell death in tumours. Therefore, the uncovering of beneficial changes in redox biology that increase CD8^+^ T cell function uncovered in this study, could allow for targeted strategies that impede tumor cell survival but at the same time enhance or maintain anti-tumor T cell function. Using T cells isolated from the skin of KDSC patients allowed us to show that a progressive loss of mitochondrial mass was associated with a loss of p38 phosphorylation and loss of pro-inflammatory cytokine production upon ex-vivo restimulation. The recovery of p38 phosphorylation and cytokine production by alterations in redox balance before restimulation suggests that there is a therapeutic opportunity in the skin to increase anti-tumor T cell function. The option to apply topical agents without the risk of systemic toxicity suggests that local administration of redox-altering agents could be an avenue to explore T cell reinvigoration in localized skin malignancies associated with a loss of p38 and proinflammatory cytokine production in TILs.

In summary, we found that fully activated CD8^+^ T_E_ cells conditioned in ↓[GLC] altered redox homeostasis in part by increased ETC CIII ROS and augmented p38 signaling, inducing increased cytotoxicity and *in vivo* persistence, and markedly improved tumor control in a mouse model of lymphoma. The redox-p38 axis could also be used to reinvigorate human KDSC CD8^+^ TILs, typically displaying exhausted or regulatory phenotypes^41^, to produce the effector cytokine IFN-γ. Taken together this suggests that this metabolic-signaling axis is a promising target to improve CD8^+^ T cell-mediated immunotherapy.

## ACKNOWLEDGEMENTS

We want to thank all past and present members of the Klein Geltink Laboratory and Dr. Bojana Rakić for helpful discussion and proofreading of the manuscript. We also want to thank Lixin Xu for expert primary cell sorting, Theresa Williams for mouse colony management, and Roger Dyer with metabolomics support.

## FUNDING

Canadian Cancer Society [Early Scholar Award to RIKG]

Natural Sciences and Engineering Research Council of Canada (NSERC), [RGPIN-2020-05390 to RIKG]

The University of British Columbia Dept of Pathology [University start-up funding to RIKG] BC Children’s Hospital Foundation [Investigator establishment funding to RIKG]

Michael Cuccione Foundation [MCF funding to RIKG]

Canadian Institutes of Health Research [CIHR; PJT-159513 to KLB] the Deutsche Forschungsgemeinschaft (SFB829, Projects B13 to M.F.)

Cologne Clinician Scientist Program (funded by the Deutsche Forschungsgemeinschaft, FI 773/15-1), Faculty of Medicine and University of Cologne (to L.B.)

UBC 4-Year Fellowship [JHO]

CIHR Canada Graduate Scholarships (CGS) awards [Doctoral to RAC] [Master’s to TS, PY and MC]

NSERC Undergraduate Student Research Awards [to TS and NS]

BC Children’s Hospital Graduate student award [ET] and Summer Studentship award [SN]

Michael Smith Health Research BC Trainee Award [ASA]

Canada Research Chair [PFL]

Michael Smith Health Research BC Scholar Award [PFL and RIKG]

## AUTHOR CONTRIBUTIONS

Conceptualization: JHO and RIKG

Experiments and data analysis: JHO, RAC, TS, SN, ET, LB, AEP, NA, JD, PY, MC, KWH, ASA, JT, TS and RIKG

Clinical specimen collection: LB

Omics analysis: PY, and YP

Funding acquisition: KLB and RIKG

Supervision: AEP, PFL, CBV, YP, MF, KLB, and RIKG

Writing - original draft: JHO, and RIKG

Writing – review & editing: all authors.

## DATA AND MATERIALS AVAILABILITY

Further information and requests for resources and reagents should be directed to and will be made available upon reasonable request by the corresponding author, Ramon I. Klein Geltink.

## STAR Methods

### Mice and adoptive T cell transfer *in vivo*

OT-I transgenic mice (C57BL/6-Tg(TcraTcrb)1100Mjb/J; Jax, 003831), CD45.1^+^ C57BL/6J (B6.SJL-Ptprca Pepcb/BoyJ; Jax, 002014), and C57BL/6J (Jax, 000664) mice were purchased from the Jackson Laboratory. CD45.2^+^ OT-I transgenic and C57BL/6J mice were housed at the Center for Molecular Medicine and Therapeutics Transgenic specific pathogen-free (SPF) mouse facility at BC Children’s Hospital Research Institute. For *in vivo* experiments, CD45.1^+^ C57BL/6J mice were maintained in the Animal Resource Centre at the BC Cancer Research Institute. All animals were cared for in compliance with the Canadian Council on Animal care and the University of British Columbia Animal Care Committee (Protocol numbers A19-024, A19-0273 and A21-0266).

For adoptive T cell transfer experiments, 12- to 14-week-old recipient CD45.1^+^ C57BL/6 female mice were housed in cohorts to match ages across experimental groups. Recipient mice were subcutaneously injected with 1 × 10^6^ E.G7-OVA lymphoma tumor cells in 100 μL serum-free RPMI1640. 8 days after tumor implantations, 3 × 10^6^ *in vitro* activated control or ↓[GLC]- conditioned OT-I CD8^+^ T cells (CD45.2^+^) were intravenously injected. 7 days after transfer, mice were bled via the saphenous vein and the cells processed for further analysis. Tumor growth was monitored every 2–3 days by measuring the average diameter of the tumor mass. Mice with tumor diameter approaching 20 mm were humanely euthanized. On day 23 after tumor implant, mice were humanely euthanized, and the tumor was harvested and further processed.

### Isolation of tumor infiltrating lymphocytes (TILs)

Tumors were excised from mice at the indicated times post-implant. Isolated tumors were mechanically chopped with a scalpel and treated with type I and type II collagenase (1 mg/mL of each, Gibco) in serum-free RPMI1640 for 40 minutes at 37°C. DNase was added (3 mg/mL, Gibco) following tissue digestion and samples were vortexed. Samples were topped up with an equal volume of RPMI1640 + 10% fetal bovine serum and filtered through a 100 μm strainer to obtain single cell suspensions and stained with antibodies as indicated below for flow cytometric analysis.

### Flow cytometric analysis of *ex vivo* cells

Blood samples were collected in EDTA containing tubes (BD) and red blood cells (RBC) were lysed with RBC lysis buffer (150 mM ammonium chloride, 10 mM potassium bicarbonate, 0.1 mM EDTA, pH 7.4), and resuspended in PBS. Spleens and tumor draining lymph nodes (left inguinal lymph node) were harvested from mice at the indicated times after transfer. Tissues were collected in PBS, dissociated with a syringe plunger, and filtered through a 70 μm strainer to obtain single cell suspensions. RBC in splenocytes were lysed with RBC lysis buffer (150 mM ammonium chloride, 10 mM potassium bicarbonate, 0.1 mM EDTA, pH 7.4). Half of the collected cells were subjected to surface proteins staining with CD8α-BV421 (1:200, clone: 53- 6.7, BioLegend), CD45.1-BV711 (1:200, clone: A20, BioLegend) and CD45.2- Alexa Flour 488 (1:200, clone: 104, BioLegend). Dead cells were excluded by staining with Fixable Viability Dye eFluor 780 (Thermo). The remainder of the collected cells were restimulated with phorbol 12- myristate 13-acetate (PMA) (50 ng/mL; Sigma) and ionomycin (500 ng/mL; Sigma) in the presence of Brefeldin A (3 µg/mL; eBioscience) and 100 U/mL rhIL-2 in TCM for 4Lhours. Cells were then subjected to intracellular protein staining for flow cytometric analysis.

### Intracellular protein detection by flow cytometry

For intracellular protein detection, αCD3/28 or PMA/Ionomycin restimulated cells were fixed and permeabilized using the Foxp3/Transcription Factor Staining Buffer Set (eBioscience), followed by anti-phospho-p38^Thr180/Tyr182^ (1:200, clone: D3F9, Cell Signaling) incubation for 1 hour at room temperature in the dark. Cells were then incubated with anti-rabbit IgG (H+L), F(ab’)2 Fragment (Alexa Fluor 647 Conjugate, 1:200, Cell Signaling) and IFN-γ-PE (1:200, clone: XMG1.2, BioLegend) for 1 hour at room temperature in the dark. MitoTracker staining was performed according to the manufacturer’s instruction (Invitrogen) using 15 nM Mitotracker Green. Samples were resuspended in 2% FBS/PBS, and acquired with a BD LSRFortessa X-20 flow cytometer. FlowJo software was used for further analysis.

### Primary cell cultures

#### Mouse T cells

CD8^+^ T cells were obtained from spleens and lymph nodes isolated from 8- to 12-week-old C57BL/6J or OT-I transgenic mice using the EasySep CD8^+^ T-cell isolation kit (Stem Cell Technologies, no. 19753) according to the manufacturer’s protocol. Within experiments, mice were age and sex matched. Isolated CD8^+^ T cells were activated using plate-bound anti-CD3 (5 µg/mL, BioXCell) and soluble anti-CD28 (0.5 µg/mL, BioXcell) in RPMI 1640 medium (Corning) supplemented with 10% fetal bovine serum (Gibco), 4 mM L-glutamine, 1% penicillin–streptomycin, and 55 µM beta-mercaptoethanol (Life Technologies) (TCM) supplemented with recombinant human (rh) IL-2 (100LU/mL, Peprotech) at 37 °C in a humidified incubator containing atmospheric oxygen supplemented with 5% CO2 for 48 hours. For OT-I cultures, single-cell suspensions of splenocytes were incubated for 48 h in medium containing rhIL-2 as above and in the presence of ovalbumin peptide SIINFEKL (100 ng/mL, Anaspec). For both WT and OT-I CD8^+^ T cell cultures, approximately 48 h after activation, medium was replaced with complete TCM medium containing 11 mM glucose and 100LU/mL rhIL-2. After a 24-hour expansion period cells were plated in fresh complete TCM media containing 100LU/mL rh IL-2, and differential glucose concentrations in the presence or absence of inhibitors or redox altering compounds as indicated in the figures and figure legends. Reagents used in primary CD8^+^ T cell culture: S3QEL-2 (10 µM, Sigma), 2,3-Dimethoxy-1,4- naphthoquinone (DMNQ; 1 µM, Sigma), SB203580 (1 µM, Sigma), Losmapimod (0.1 µM, Selleck Chemicals), N-acetyl-L-cysteine (NAC; 10 mM, Sigma), 6-Aminonicotinamide (6-AN; 250LμM, Sigma).

#### Human healthy control CD8^+^ T cells

Naïve CD8^+^ T cells were isolated using a human naïve CD8+ T cell isolation kit (Stemcell Technologies) from peripheral blood mononuclear cells obtained by Ficoll paque density gradients out of blood donated by healthy individuals as approved by the ethics review board at UBC and BCCHR, protocol number H19-02734. Following isolation, T cells were resuspended at 1 x 10^6^ cells per ml and activated using anti-CD2/3/28 tetramers (Stemcell Technologies) for 3 days in proprietary serum free ImmunoCult media (Stem Cell Technologies) containing 200 U/ml rhIL-2 (Peprotech). At day 3, fresh media containing IL2 was added for a further 48 hours. For experiments, cultures were split into 10mM glucose containing ImmunoCult (control), or glucose free ImmunoCult media supplemented with 1mM glucose (↓[GLC]) for 20 hours before experimental analysis.

#### Human Tumor Infiltrating Lymphocytes

Human TILs were isolated from 4 mm punch biopsies from patients with KDSC who underwent surgery for their tumors at the Department of Dermatology and Venereology at the University of Cologne. The study was conducted according to the principles expressed in the Declaration of Helsinki and approved by the local Ethic Committee (*Ethikkommission*) of the University of Cologne (votes #08-144, #20-1082 and #19-1146). All donors provided written informed consent for the collection of blood and subsequent analyses.

For *ex vivo* staining, biopsies were collected in PBS and dissociated using the Whole Skin Dissociation Kit (Miltenyi). For MitoTracker staining, biopsies of the tumor center, tumor margin, and adjacent skin of n=15 samples (n=6 basal cell carcinoma (BCC), n=8 squamous skin cell carcinoma (SCC), n=1 Morbus Bowen) were incubated with the Whole Skin Dissociation Kit for 3hs according to the manufacturer’s instructions. Then, MitoTracker staining was performed according to the manufacturer’s protocol (Invitrogen) using 15 nM Mitotracker Green (incubation for 20 minutes at 37°C). Surface protein staining was performed with Alexa Fluor® 700 anti-human CD3 (1:400, clone: HIT3a, BioLegend) and Brilliant Violet 605™ anti-human CD8 (1:400, clone; SK1, BioLegend) in FACS buffer for 20 minutes at 4°C. Dead cells were excluded by staining with Fixable Viability Dye eFluor 780 (Thermo).

To assess p-p38 and IFN-γ expression in human TILs *ex vivo*, 4mm punch biopsies from tumor margin and adjacent skin from n=4 SCC samples were incubated with the Whole Skin Dissociation Kit overnight. Next, cells were restimulated with αCD2/3/28 tetramers (STEMCELL Technologies) in RPMI containing 10% FCS and 200U/ml rhIL-2 for 5 hours. Brefeldin A (ThermoFisher Scientific) was added for the final 2.5 hours. Surface staining was performed as described above. Restimulated cells were fixed and permeabilized using the Foxp3/Transcription Factor Staining Buffer Set (eBioscience), followed by anti-phospho- p38Thr180/Tyr182 (1:200, clone: D3F9, Cell Signaling) incubation for 1 hour at room temperature in the dark. Cells were then incubated with anti-rabbit IgG (H+L), (ab’)2 Fragment (Alexa Fluor 647 Conjugate, 1:200, Cell Signaling) and PE/Cy7 anti-human IFN-γ (1:400, clone: B27, BioLegend) for 30 minutes at 4°C.

For DMNQ treatments, TIL cultures were generated out of 4 mm punch biopsies from tumor margins of n=2 SCC and n=2 BCC. Tissue pieces were placed into 48-well plates and incubated with 1 mL enriched medium (RPMI 1640 with L-glutamine, 1% sodium pyruvate, 1% penicillin/streptomycin, and 55 μmol/L 2-mercaptoethanol), and 10% human serum plus 10U/mL IL-2 at 37 °C, 5% CO2. After two days, 10 U/mL fresh IL-2 were added. On day 4, the tissue piece was removed, and the migrated T cells were collected and expanded with ImmunoCult™ Human αCD2/3/28 tetramers for up to 36 hours in T-cell expansion media containing IL 2 according to the manufacturer’s protocol. T-cell enriched medium supplemented with IL-2 was replaced every third to fourth day. DMNQ was added to cultures for 24 hours, followed by restimulation with αCD2/3/28 tetramers overnight followed by staining for the indicated markers. For flow cytometry acquisition, cells were resuspended in FACS buffer and put on ice. Acquisition of cells was performed on an Attune NxT Flow Cytometer (Thermo Fisher Scientific). Data were analyzed using FlowJo software (Tree Star, Ashland OR).

### Cell lines

The mouse E.G7 lymphoblast cell line expressing OVA (E.G7-OVA) was purchased from the American Type Culture Collection (ATCC no. CRL-2113; RRID: CVCL_3505). Cells were cultured in TCM supplemented with 10% fetal bovine serum (Gibco), 1% penicillin– streptomycin (Hyclone). The mouse B16F10-OVA melanoma cell line was a kind gift from D. Zehn. Cells were maintained in DMEM (B16F10-OVA) supplemented with 10% fetal bovine serum (Hyclone), 1% penicillin–streptomycin (Hyclone). All cells were maintained at 37 °C in a humidified incubator containing atmospheric oxygen supplemented with 5% CO_2_. Cells were routinely tested for mycoplasma contamination using an EZ-PCR™ Mycoplasma Detection Kit (Biological Industries Israel Beit-Haemek Ltd.) in accordance with the manufacturer’s protocol.

### Retrovirus production and retroviral gene transduction

Short hairpin RNA (shRNA) (mouse *Mapk14*, TRCN0000023119) oligonucleotides containing 5’ AgeI and 3’ EcoRI 4 nucleotide overhangs were synthesized by IDT and annealed in 1 × cutsmart buffer (NEB) followed by phosphorylation with T4 Polynucleotide Kinase (NEB) and cloned into the AgeI-EcoRI pMKO.1-GFP retroviral vector (pMKO.1 GFP was a gift from William Hahn (Addgene plasmid # 10676 ; http://n2t.net/addgene:10676 ; RRID:Addgene_10676)). pUC57-TPNOX was a gift from Vamsi Mootha (Addgene plasmid # 87853 ; http://n2t.net/addgene:87853 ; RRID:Addgene_87853). MluI was used to transfer the TPNOX coding sequence to an MSCV-I-GFP retroviral vector. To prepare retroviral particles, 3 × 10^6^ logarithmically expanding HEK293T cells were plated on 10-cm culture dishes and transfected with 10 µg pMKO.1-*Mapk14* shRNA-GFP or MSCV-TPNOX-FLAG-IRES-GFP combined with 6 µg pCL-eco using calcium phosphate precipitation (pCL-Eco was a gift from Inder Verma (Addgene plasmid # 12371 ; http://n2t.net/addgene:12371 ; RRID:Addgene_12371). 16 hours after transfection, media was replaced with 9 mL TCM. Retroviral supernatants in TCM were collected ∼48, 60 and 72 hours after transfection and filtered through 0.2 µm filters. For transduction of T cells, 48 hours post αCD3/28 activation as above, 2 × 10^6^ fully-activated CD8^+^ T cells were resuspended in 5 mL *Mapk14* shRNA-GFP or TPNOX-FLAG-GFP retrovirus supernatants supplemented with rhIL-2 (∼MOI of 3) and centrifuged at 30°C for 90 minutes at 2,000 rpm in the presence of polybrene (8 μg/mL, Sigma), followed by 90 minutes recovery at 37°C in a humidified incubator. Next, half of the media was replaced, and cells cultured in complete TCM containing rhIL-2. Media was replaced 24 hours later, and GFP-positive cells sorted 48 hours after transduction for further analysis.

### Co-culture of OT-I T_E_ cells and OVA-expressing tumor cell lines

50,000 B16F10-OVA cells were plated in duplicate wells of a flat bottom 96-well plate. After 24 hours, tumor cell media was replaced with fresh TCM containing 200,000 control or glucose ↓[GLC]-conditioned T_E_ cells. Brefeldin A was added 20 hours after the start of co-culture for 4 hours followed by cell surface staining with CD8α-BV421 (1:400, clone: 53-6.7, BioLegend) for 30 minutes at room temperature in the dark; Fixable Viability Dye eFluor 780 (1:10,000, 65- 0865-18, Thermo) was used to exclude dead cells. Intracellular phospho-p38^Thr180/Tyr182^ and IFN- γ were measured as described above.

### Redox-sensitive green fluorescent protein (roGFP) sensor analysis

CD8^+^ T cells containing MSCV retrovirus expressing Cyto-roGFP and mitomatrix-roGFP were generated and packaged as previously described ^18^. roGFP-expressing control or ↓[GLC]- conditioned T_E_ cells were restimulated with anti-CD3/28 antibodies or Dynabead Mouse T-Activator CD3/28 (Thermo) for 4 hours in glucose sufficient (10 mM) media, and subjected to flow cytometric analysis immediately after restimulation. Excitation with a 488- and 405-nm laser was used to quantify reduced and oxidized signal, respectively on a BD LSRFortessa X20 flow cytometer. FlowJo software was used to generate the derived parameter of the two fluorescent signals.

### *In vitro* tumor killing assay

E.G7-OVA were co-cultured with OT-I T_E_ cells at 2:1 or 1:1 or 1:2 ratio for 24 hours in sterile 96-well v-bottom plates. Samples were then stained with CD8α-APC (1:300, clone: 53-6.7, BioLegend) and 0.2 μg/mL propridium iodide (PI). E.G7-OVA (CD8 ^neg^PI^+^) and CD8^+^ T cell (CD8α^+^PI^+^) death was measured by flow cytometry on a BD LSRFortessa X-20 flow cytometer.

### mRNA expression analysis by real-time quantitative PCR (qRT-PCR)

Total RNA was extracted using TRIzol reagent (Invitrogen) and reverse transcribed into cDNA using a High-Capacity cDNA Reverse Transcription Kit (Applied Biosystems) in accordance with the manufacturer’s protocol. Specific primers for genes (*Atf4, Ddit3, Ifng* and, *Prf1*) and *Hprt* (internal control) were purchased from IDT. qPCR was performed in triplicate with PowerTrack™ SYBR Green Master Mix (Applied Biosystems) using a ViiA 7 Real-Time PCR System (Thermo). The thermal cycling conditions were 2Lminutes at 95°C, followed by 40 cycles of 95 °C for 5Lseconds and 60 °C for 30 seconds and one cycle of 95 °C for 15 seconds, 60 °C for 1 minute, and 95 °C for 15 s. Results were analyzed by the 2^−ΔΔCt^ method and normalized to *Hprt* expression. qRT-PCR primers:

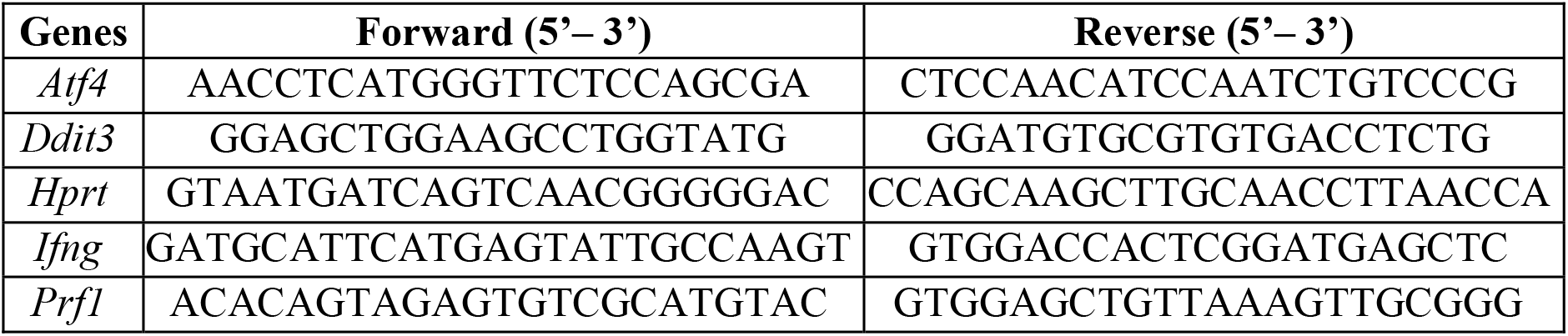

### Immunoblotting

Cells were washed with ice-cold PBS and lysed in lysis buffer containing 20 mM Tris-HCl (pH 7.5), 150 mM NaCl, 1LmM Na_2_EDTA, 1 mM EGTA, 1% Triton X-100, 2.5LmM sodium pyrophosphate, 1 mM beta-glycerophosphate, 1LmM Na_3_VO_4_ and 1 μg/mL leupeptin (Cell Signaling) supplemented with 1 mM phenylmethyl sulfonyl fluoride (PMSF, Sigma). Samples were frozen and thawed three times, followed by centrifugation at 14,000 rpm for 10Lminutes at 4L°C. Cleared protein lysate was denatured with 4 × LDS loading buffer and 50 mM DTT for 10Lmin at 70 °C and loaded on precast Bolt or NuPAGE 4–12% Bis–Tris Plus gels (Thermo). Proteins were transferred onto nitrocellulose membranes (Invitrogen) using the iBlot 2 dry blotting system (Life Technologies, 8.5 minutes at 30V) according to the manufacturer’s protocols. Membranes were blocked with 5% w/v non-fat dry milk (Bio-rad) and 0.1% Tween-20 (Sigma) in Tris-buffered saline (TBS-T) and incubated with the appropriate antibodies in 5% w/v BSA (BioShop) in TBS (pH 7.6) with 0.1% Tween-20 at 4 °C overnight (primary antibodies were added at a 1:2,000 dilution and from Cell Signaling Technology unless otherwise indicated). The antibodies used were: anti-phospho-p38^Thr180/Tyr182^ (cat. 4511, 1:2,000); anti-p38 (total) (cat. 9218, 1:2,000); anti-phospho-ATF-2^Thr71^/ ATF-7^Thr53^(cat. 15441, 1:2,000); anti-ATF-2/ATF-7 (total) (cat. 82870; 1:2,000); anti-phospho-4E-BP1^Thr37/45^ (cat. 2855; 1:4,000); anti-4E-BP1 (cat. 9644; 1:4,000); anti-ATF4 (cat. 11815; 1:2,000); anti-CHOP (cat. 2895; 1:2,000); and anti-α-Tubulin (Sigma, T6199; 1:10,000). All primary antibody incubations were followed by incubation with secondary HRP-conjugated anti-rabbit (Invitrogen, cat. 31460) or anti-mouse antibodies (Invitrogen, cat. 31430) in 5% milk and 0.1% Tween-20 in TBS (1:15,000) and visualized on radiosensitive film (Diamed) using chemiluminescent substrate (SuperSignal West Pico or Femto, Thermo).

### Protein oxidation detection (OxyBlot)

Protein carbonylation, as a readout of protein oxidation, was measured using an OxyBlot Protein oxidation Detection kit (Sigma) according to the manufacturer’s recommendations. Total protein lysates were denatured in 6% sodium dodecyl sulfate (SDS). Carbonyl groups in the protein side chains were derivatized to 2, 4-dinitrophenylhydrazine (DNP) by DNP hydrazine. Prepared samples were separated on precast Bolt 4-12% Bis-Tris Plus gels (Thermo) transferred to nitrocellulose membranes (Invitrogen) using an iBLOT2 (Life Technologies, 8.5 minutes at 30V) after blocking with 5% milk in TBS-tween the membranes were incubated with anti-DNP primary antibody (1:150) overnight at 4 °C, followed by HRP-conjugated secondary antibody (1:300).

### Seahorse metabolic flux analysis

Extracellular acidification rate (ECAR) and oxygen consumption rate (OCR) were measured using using a XFe 96 Extracellular Flux Analyzer (Agilent). 2 × 10^5^ mouse or human T cells per well (≥ 3 wells per sample) were spun onto poly-D-lysine coated seahorse 96-well plates in 40 µL of unbuffered RPMI media, topped up to 180 uL and preincubated at 37°C for a minimum of 45 minutes in the absence of CO_2_. For control T_E_ cells, unbuffered RPMI media containing 10 mM glucose, 2 mM glutamine, and 1 mM sodium pyruvate was used. For ↓[GLC]-conditioned T_E_ cells, unbuffered RPMI media containing 1 mM glucose, 2 mM glutamine, and 1 mM sodium pyruvate was used. Mitochondrial phenotypes were assessed using the mitochondrial stress test by consecutive injection of: 1 μM oligomycin, 1.5 μM flurorcarbonyl cynade phenylhydrazon (FCCP), 100 nM rotenone + 1 μM antimycin A, and 50 mM 2-deoxy-D-glucose (2-DG) (all Sigma) as indicated.

### Measurement of extracellular metabolites

Extracellular concentrations of glucose, lactate, glutamine and glutamate were measured in the T cell culture media supernatants before experiments were performed on control and ↓[GLC]- conditioned T_E_ cells. Briefly, ∼500 µL media was collected after overnight culture followed by centrifugation for 400 x g for 5 minutes at 4°C to remove cells. Quantification was performed using a Roche Cedex Bio analyzer using colorimetric assays following manufacturer’s protocols, and measurements were corrected by comparison to media from wells not containing cells incubated over the same time frame.

### NADPH/NADP^+^-glo assay

250,000 control or ↓[GLC]-conditioned T_E_ cells were washed once in ice cold PBS and lysed in 100 μL of lysis buffer (1% dodecyltrimethylammonium bromide (DTAB):PBS = 1:1). The NADPH/NADP^+^ ratio was measured using an NADPH/NADP^+^ glo Assay Kit (Promega) according to the manufacturer’s instructions. For NADPH measurement, 50 μL of lysate was transferred to PCR tubes and incubated at 60°C for 15 minutes. For NADP^+^ measurement, 50 μL of lysate was transferred to PCR tubes containing 25 μL of 0.4 N HCl, and incubated at 60°C for 15 minutes. After incubations, samples were allowed to equilibrate to room temperature for 10 minutes and then neutralized with 25 μL of 0.5 M Tris (for NADPH) or 50 μL of 0.4 N HCl (for NADP^+^). Prepared samples were mixed with NADPH/NADP^+^-glo detection reagent in a white 384-well luminometer plate and incubated at room temperature for 45 minutes. Luminescence was measured using an Infinite® 200 plate reader (Tecan).

### Metabolomics by LC-MS

Cells were washed with ice cold 150 mM ammonium formate buffer, 50% −20°C HPLC-grade methanol was added and samples kept on dry ice, followed by a phase-separation using acetonitrile, water, and dichloromethane after vigorous vortexing for 60 seconds per sample. The aqueous phase was collected by pipette and stored at −80 before shipping to the University of Ottawa Metabolomics Center. Samples were dried in a cold trap overnight at −1°C. Pellets were resuspended in HPLC water and injected into an Agilent 6540 UHD Accurate-Mass Q-TOF LC/MS system coupled to ultra-high pressure liquid chromatography (UHPLC, 1290 Infinity LC System). Metabolite masses and spectra were analyzed using Masshunter software.

### Single cell RNA-seq analysis

Single-cell RNA-seq data of control and ↓[GLC]-conditioned T_E_ cells GSE152018 ^18^ was compared to that of B16 murine melanoma tumor infiltrating CD8^+^ T cells GSE116390 ^38^. The datasets were integrated using Seurat v3 ^62^ for joint analysis. The original labels introduced in the CD8^+^ TIL dataset were used to label the T cell subsets. The resulting dataset contained a total of 9458 cells (5884 and 3574 from the in vitro T_E_ cells and the CD8^+^ TIL datasets, respectively). 1967 highly variable genes were identified and used for dimensionality reduction using Principal Component Analysis (PCA). For visualization, t-SNE was run using the first 10 principal components, perplexity = 30, and seed set to 123. For clustering, shared nearest neighbor modularity optimization clustering algorithm from Seurat v3 was used with resolution parameter = 0.5. A total of 7 clusters were obtained. Marker genes of cluster 3 with an adjusted P value < 0.05 were identified using FindMarkers implemented in Seurat v3. Gene set enrichment analysis was performed against KEGG gene pathways. Statistically significant pathways with P value < 0.05 were identified using a hypergeometric test.

### Statistical analysis

Statistical analysis was performed using Prism 9 software (version 9.5.0; Graphpad) and results are represented as mean ± SEM, unless otherwise indicated in the figure legends. Comparisons for two groups were calculated using unpaired two-tailed Student’s *t-*tests, comparisons of more than two groups were calculated using one-way ANOVA with Tukey’s multiple comparison tests. Comparisons of over time were calculated using two-way ANOVA followed Šídák’s multiple comparisons test

