## Supplementary material for "Inducing an oxidized redox-balance improves anti-tumor CD8^+^ T cell function": Figure S1

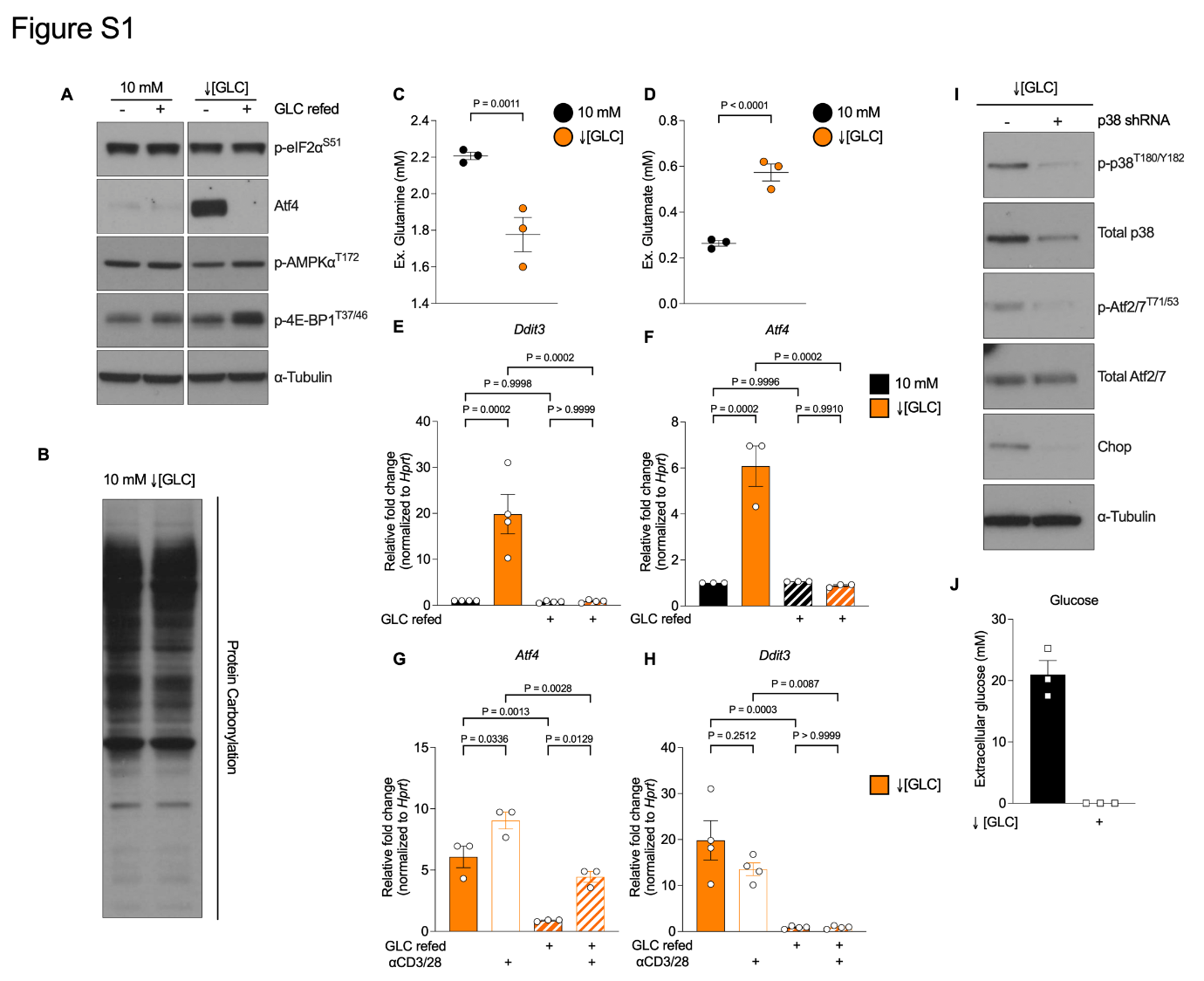

**Figure S1.** **↓[GLC]** **conditioning activates p38 but not canonical ISR signaling in of CD8^+^ T cells**. (**A**) Representative immunoblot analysis of protein extracts isolated from control and ↓[GLC]-conditioned T_E_ cells with or without a 4-hour glucose refeed, probed for phosphorylated eIF2α at S51 (p-eIF2α^S51^), Atf4, phosphorylated AMPKα1 at T172 (p-AMPKα^T172^), phosphorylated 4E-BP1 at T37/46 (p-4E-BP1^T37/46^). α-Tubulin was used as a loading control. Data are representative of n > 6 independent experiments with 3 biological replicates each. (**B**) Representative OxyBlot analysis of protein extracts from control and ↓[GLC]-conditioned T_E_ cells, probed for 2, 4-dinitrophenylhydrazine (DNP) to detect protein carbonylation. n = 3 biological independent samples. Quantification of extracellular (Ex.) (**C**) glutamine and (**D**) glutamate in the medium of T_E_ cells cultured after 20 hours with or without ↓[GLC] conditioning. n = 3 biological replicates. qRT-PCR analysis of (**E**) *Ddit3* and (**F**) *Atf4* 4 hours after glucose (GLC, 10 mM) refeeding of control and ↓[GLC]-conditioned T_E_ cells. n = 3 biological replicates are shown. qRT-PCR analysis of (**G**) *Atf4* and (**H**) *Ddit3* 4 hours after αCD3/28 restimulation with or without GLC refeeding of ↓[GLC]-conditioned T_E_ cells. n = 3 biological replicates are shown. (**I**) Representative immunoblot analysis of p38 pathway in ↓[GLC]-conditioned T_E_ cells transduced with or without p38 (*Mapk14*) shRNA. Data are representative of 3 biological replicates. (**J**) Quantification of extracellular glucose in the medium of fully activated human T cells culture after 20 hours with or without ↓[GLC] conditioning. n = 3 biological independent samples are shown. Data are presented as mean ± SEM and *P* values are determined by two-tailed Student’s *t*-test or one-way ANOVA with multiple comparisons.
