## Supplementary material for "Inducing an oxidized redox-balance improves anti-tumor CD8^+^ T cell function": Figure S2

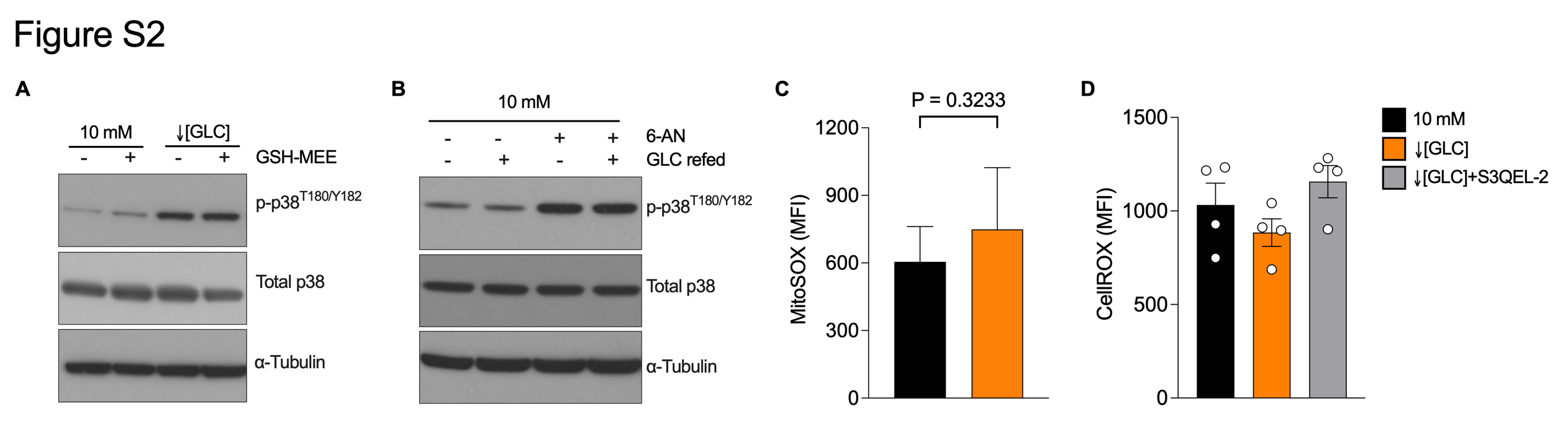

**Figure S2. Antioxidants did not blunt p38 activity in T_E_ cells.** (**A**) Representative immunoblot analysis of p-p38^T180/Y182^ and total p38 in control and ↓[GLC]-conditioned T_E_ cells with or without cell-permeable reduced glutathione ethyl ester (GSH-MEE, 4 mM) for 20 hours. (**B**) Representative immunoblot analysis of p-p38^T180/Y182^ and total p38 in control T_E_ cells with or without 6-Aminonicotinamide (6-AN, 250 μM) for 4 hours. (**A-B**) α-Tubulin was used as a loading control. Data are representative of n = 3 biological replicates each. (**C**) Quantification and representative flowcytometric contour plots of mitoSOX staining (**D**) Mean fluorescence intensity (MFI) of CellROX staining as a readout of cellular reactive oxygen species (ROS) in control and ↓[GLC]-conditioned T_E_ cells treated with vehicle or S3QEL-2 (10 μM). Data are presented as mean ± SEM. No significant differences were identified by one-way ANOVA with multiple comparisons.
