## Supplementary material for "Inducing an oxidized redox-balance improves anti-tumor CD8^+^ T cell function": Figure S3

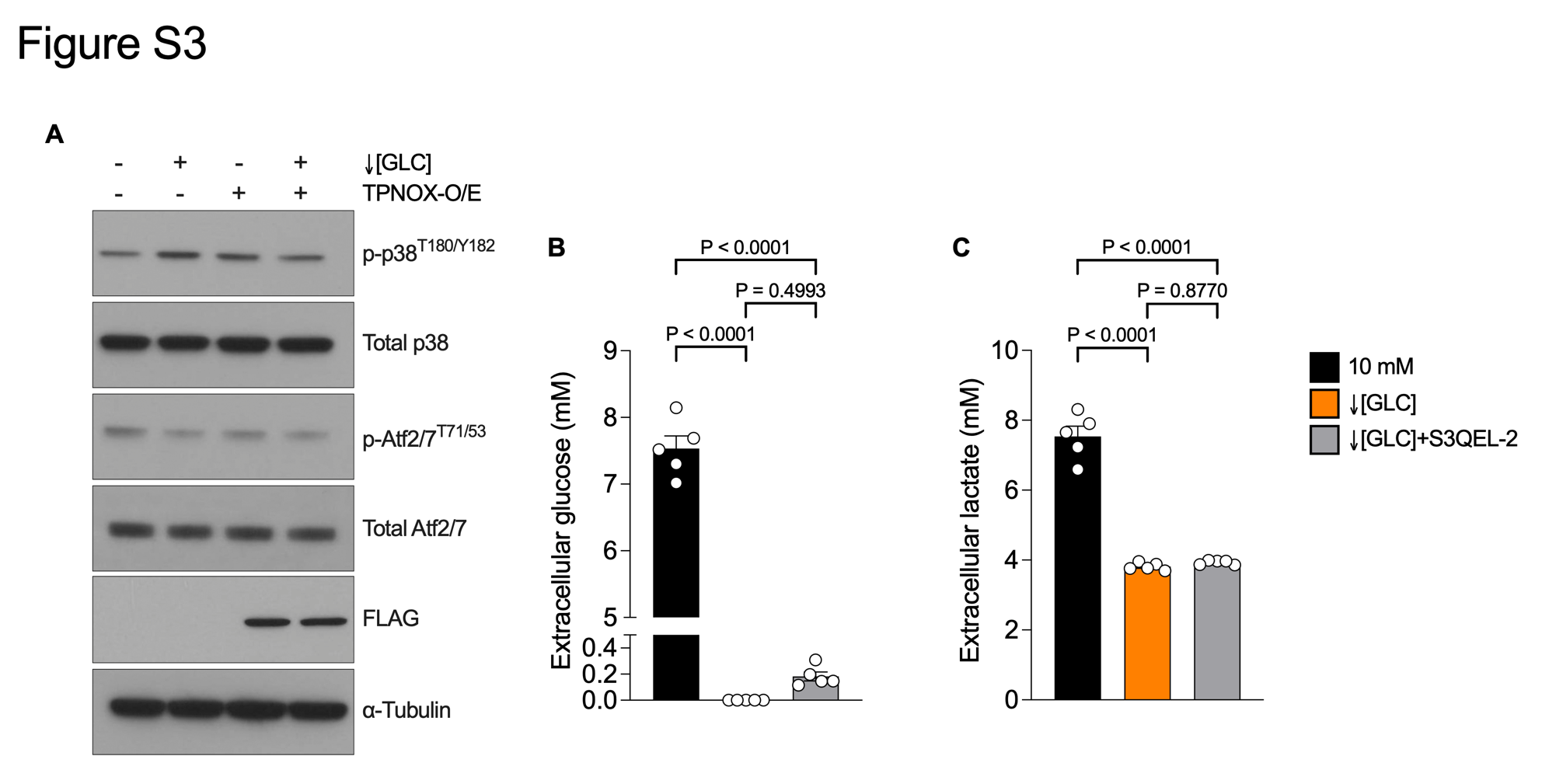

**Figure S3. The effects of redox modulators on p38 activation and glucose uptake.** (**A**) Representative immunoblot analysis of p38 pathway in control and↓[GLC]-conditioned T_E_ cells transduced with or without flag-tagged TPNOX overexpression (O/E). Flag was probed as a transduction control and α-Tubulin was used as a loading control. Data are representative of 3 biological replicates. Quantification of extracellular (**B**) glucose and (**C**) lactate in the medium of control and ↓[GLC]-conditioned T_E_ cells cultured with or without S3QEL-2 (10 μM) for 20 hours. n = 5 biological replicates are shown. Data are presented as mean ± SEM and *P* values are determined by one-way ANOVA with multiple comparisons.
