## Supplementary material for "Inducing an oxidized redox-balance improves anti-tumor CD8^+^ T cell function": Figure S4

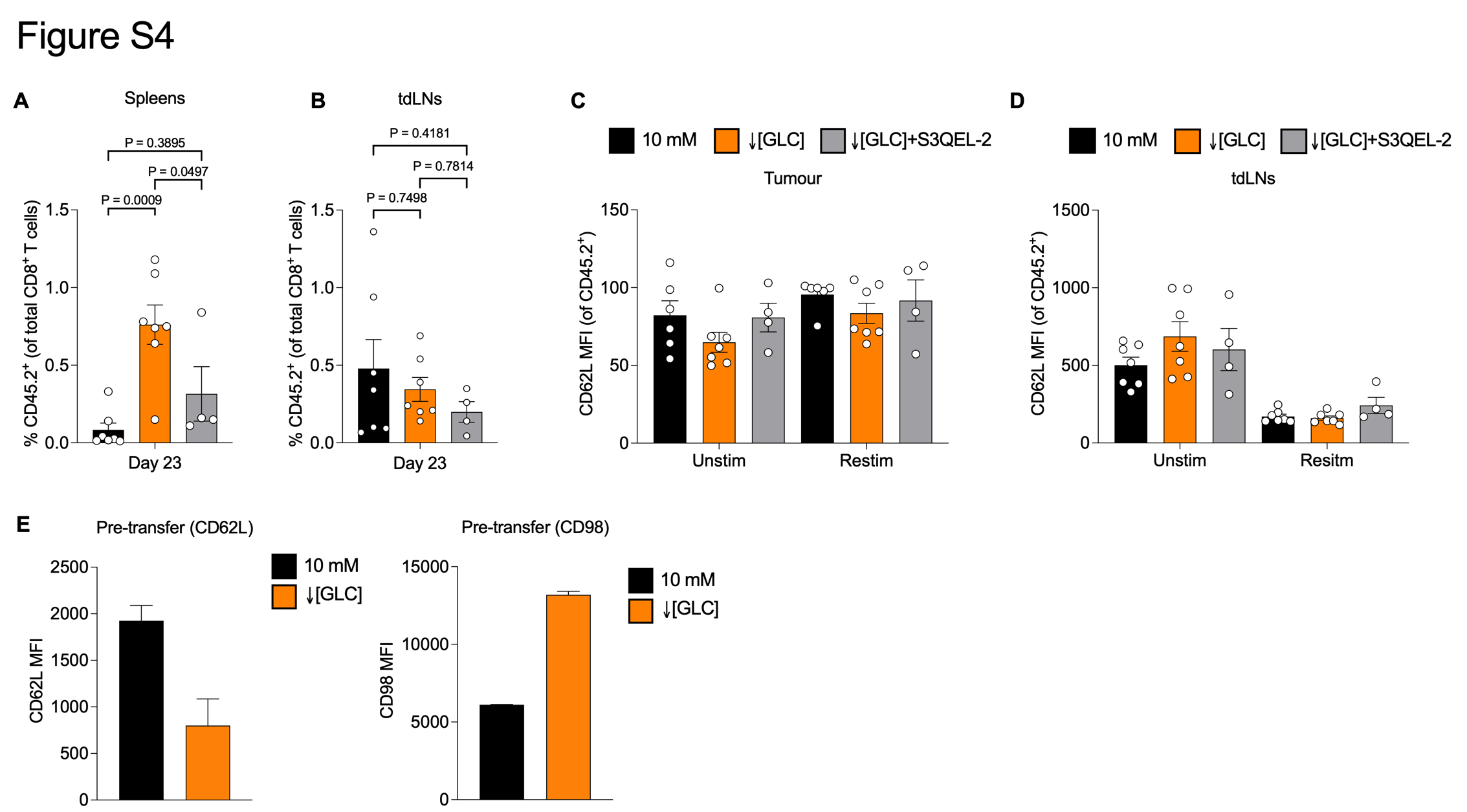

**Figure S4. Distinct immunological phenotypes of ↓[GLC]-conditioned T_E_ cells.** (**A**) Spleens and (**B**) tumor draining lymph nodes (tdLNs) were harvested on day 23 post E.G7-OVA implant as described in Figure 4a, and the frequency of CD45.2^+^ donor CD8^+^ T cells was measured by flowcytometry. (**C**) Tumor and (**D**) tdLNs were harvested on day 23 post E.G7-OVA implant as described in Figure 4a, and the mean fluorescence intensity (MFI) of CD62L was measured after *ex vivo* restimulation with PMA/ionomycin for 4 hours. (**E**) Quantification of CD62L and CD98 MFI in control and ↓[GLC]-conditioned T_E_ cells before adoptive cell transfer. Data are presented as mean ± SEM and *P* values are determined by one-way ANOVA with multiple comparisons.
