## Supplementary material for "Inducing an oxidized redox-balance improves anti-tumor CD8^+^ T cell function": Figure S5

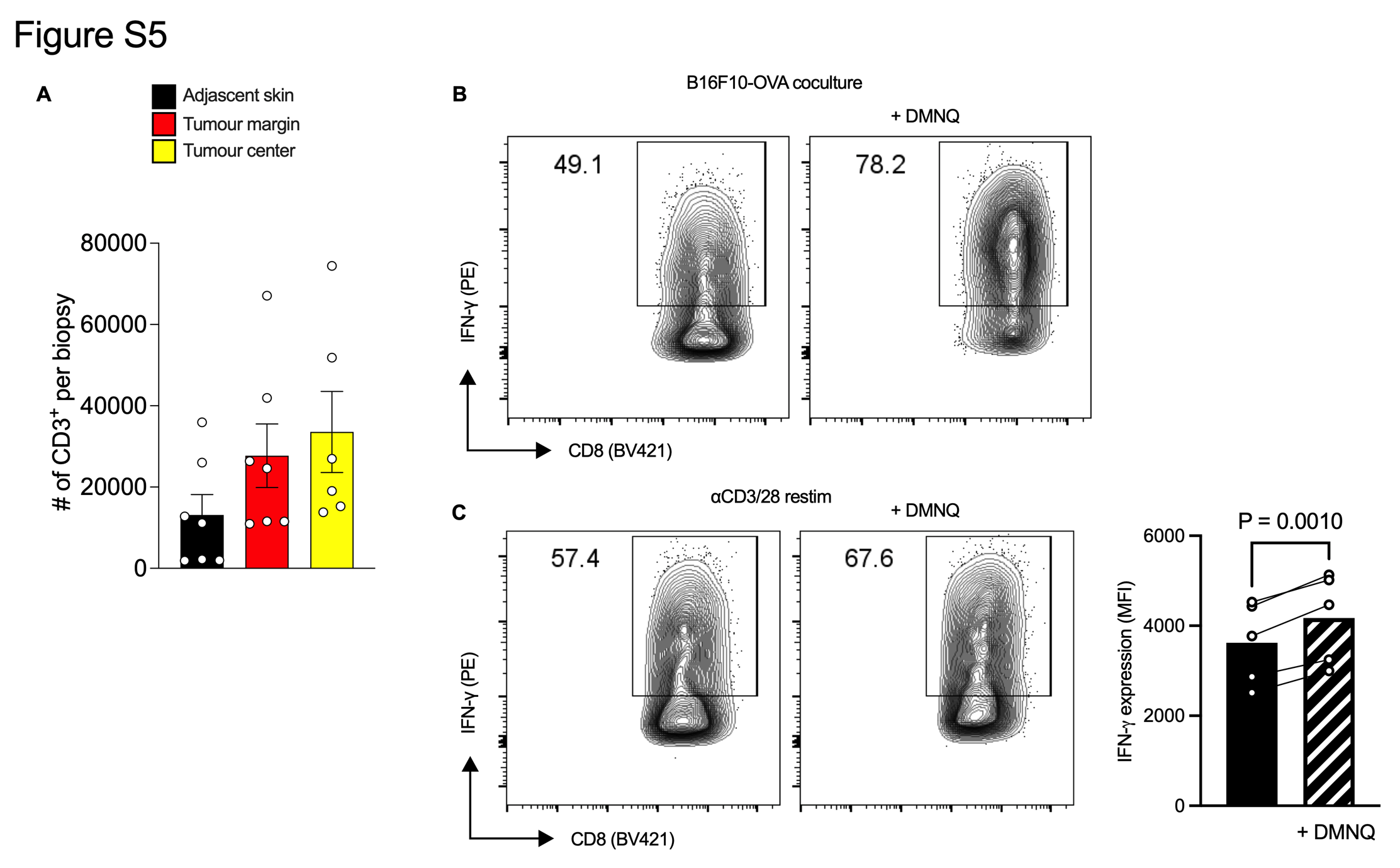

**Figure S5. Induction of redox cycling in CD8^+^ T_E_ increases IFN-γ production.** (**A**) The number of human CD3^+^ tumor infiltrating T cells from punch biopsies of adjacent skin, tumor-margin or -center was assessed by flow cytometry. Representative flow contour plots of IFN-γ measurement in mouse T_E_ cells that were (**B**) cocultured with B16F10-OVA or (**C**) restimulated with αCD3/28 in the absence or presence of DMNQ for 24 hours. *P* values are determined by paired T test, quantifying two independent experiments with n=5 biological replicates total.
